# *Oecophyllibacter saccharovorans* gen. nov. sp. nov., a bacterial symbiont of the weaver ant *Oecophylla smaragdina*

**DOI:** 10.1101/2020.02.15.950782

**Authors:** Kah-Ooi Chua, Wah-Seng See-Too, Jia-Yi Tan, Sze-Looi Song, Hoi-Sen Yong, Wai-Fong Yin, Kok-Gan Chan

## Abstract

In this study, bacterial strains Ha5^T^, Ta1 and Jb2 were isolated from different colonies of weaver ant *Oecophylla smaragdina*. They were distinguished as different strains based on matrix-assisted laser desorption ionization-time of flight (MALDI-TOF) mass spectrometry and distinctive random-amplified polymorphic DNA (RAPD) fingerprints. Cells of these bacterial strains were Gram-negative, rod-shaped, aerobic, non-motile, catalase-positive and oxidase-negative. They were able to grow at 15–37°C (optimum, 28–30°C) and in the presence of 0–1.5 % (w/v) NaCl (optimum 0%). Their predominant cellular fatty acids were C_18:1_ *ω*7*c*, C_16:0_, C_19:0_ *ω*8*c* cyclo, C_14:0_ and C_16:0_ 2-OH. Strains Ha5^T^, Ta1 and Jb2 shared highest 16S rRNA gene sequence similarity (94.56–94.63%) with *Neokomagataea tanensis* NBRC106556^T^ but were phylogenetically closer to *Bombella* spp. and *Saccharibacter floricola* DSM15669^T^. Both 16S rRNA gene sequence-based phylogenetic analysis and core gene-based phylogenomic analysis placed them in a distinct lineage in family *Acetobacteraceae*. These bacterial strains shared higher than species level thresholds in multiple overall genome-relatedness indices which indicated that they belonged to the same species. In addition, they did not belong to any of the current taxa of *Acetobacteraceae* as they had low pairwise average nucleotide identity (≤70.7%), *in silico* DNA-DNA hybridization (≤39.5%) and average amino acid identity (≤66.ü%) values with all the type members of the family. Based on these results, bacterial strains Ha5^T^, Ta1 and Jb2 represent a novel species of a novel genus in family *Acetobacteraceae,* for which we propose the name *Oecophyllibacter saccharovorans* gen. nov. sp. nov., and strain Ha5^T^ as the type strain.

## Introduction

The family *Acetobacteraceae* within the order Rhodospirillales consists of Gram-negative, aerobic and non-spore forming bacteria (Gillis and De Ley, 1980, Yamada, 2016). Members of this family are known for their oxidative capability on carbohydrates and a major group of them are recognized as acetic acid bacteria (AAB) (Yamada and Yukphan, 2008). The inability of AAB to completely oxidize alcohols and other carbohydrates leads to accumulation of partially oxidized metabolic products such as aldehydes, ketones and other organic acids in the growth medium (Kersters *et al.*, 2006). Their great metabolic potentials make them industrially important for manufacture of vinegar, foods and different chemical compounds (Gomes *et al.*, 2018).

The members of family *Acetobacteraceae* are ubiquitous in the environments. At the time of writing, family *Acetobacteraceae* contains 45 validly published genera (two synonyms) (http://www.bacterio.net/Acetobacteraceae.html). Although species of these genera were mostly isolated from sugary, alcoholic or acidic habitats including fermented food and alcoholic beverages (Cleenwerck *et al.*, 2007, Kim *et al.*, 2018, Spitaels *et al.*, 2014), a number of them are insect-associated (Crotti *et al.*, 2010). Species of genera *Gluconobacter, Acetobacter* and *Asaia* have been cultivated from fruit fly *Drosophila melanogaster, Bactrocera oleae* and mosquito *Anopheles stephensi* respectively (Favia *et al.*, 2007, Kounatidis *et al.*, 2009, Roh *et al.*, 2008). Recently, a new genus *Bombella* had been reported with two novel species isolated from the digestive tracts of multiple bee species (Li *et al.*, 2015, Yun *et al.*, 2017).

It is believed that symbiotic interaction with bacteria is one of the key attributes to remarkable adaptability and thus the evolutionary success of insects. For insects that survive on protein-poor diets such as honeydew and plant sap, the symbiotic bacteria provide amino acids as nutritional support and contribute to fitness of the host (Feldhaar *et al.*, 2007). The use of high-throughput sequencing technology revealed a diverse group of *Acetobacteraceae* taxa constituting insect-associated microbial community including ants (Brown and Wernegreen, 2019), bees (Crotti *et al.*, 2013) and fruit flies (Kounatidis *et al.*, 2009, Yong *et al.*, 2019). These *Acetobacteraceae* taxa are known to survive acidic environment in insect guts and tolerate sugar-rich diets of the host (Crotti *et al.*, 2016).

The weaver ant *Oecophylla smaragdina* (Fabricius, 1775) is an abundant obligate arboreal ant species in Southeast Asia and northern Australia (Pimid *et al.*, 2014). The ability of *O. smaragdina* to build nest directly from leaves still attached on host trees enables it to survive on a large number of host plant species. In addition to feeding on sugar-rich plant exudates such as extrafloral nectar and honeydews, the highly aggressive major workers of *O. smaragdina* prey on a wide range of intruding arthropods including insect pests (Holldobler and Wilson, 1990). Their territorial behaviour has been exploited in biological control of many insect pests on tropical tree crops such as mango (Peng and Christian, 2004), cocoa (Way and Khoo, 1989) and cashew (Peng *et al.*, 1995). While its role as biological control agent continues to expand, the study about microbiota of *O. smaragdina* is still limited.

By 16S rRNA gene amplicon sequencing, our previous study has shown that several operational taxonomic units (OTUs) of the family *Acetobacteraceae* were consistently detected in microbiota of *O. smaragdina* (Chua *et al.*, 2018). These OTUs were assigned to genus *Neokomagataea* and were categorized as core members of the *O. smaragdina* microbiome. In this study, we attempt to cultivate the *Acetobacteraceae* taxa from *O. smaragdina* and report the taxonomic characterizations of strains Ha5^T^, Ta1 and Jb2 using polyphasic approaches. Based on the results obtained, we describe the three bacterial strains as a novel species of a novel genus in the family *Acetobacteraceae*.

## Materials and methods

### Ant specimen collection, isolation of *Acetobacteraceae* and culture conditions

Major workers of *O. smaragdina* were caught alive from the field in 3 distant locations of Kuala Lumpur and Petaling Jaya, Malaysia (Supplementary data Table S1) during June and July 2017 and promptly transferred in sterile container to laboratory in the University of Malaya (Kuala Lumpur, Malaysia). The ants were cold-anesthetized at 4°C for 5 min, surface sterilized with 70% v/v ethanol for 1 min and rinsed with sterile distilled water. The gasters of 5 ants were detached and homogenized in 500 μL 1× phosphate-buffered saline (PBS) with a sterile micro-pestle (Lin *et al.*, 2016).

Different pre-enrichment media were used for isolation of *Acetobacteraceae* from the homogenates. Enrichment medium A was composed of 2.0% (w/v) D-glucose, 0.5% (v/v) ethanol, 0.5% (w/v) peptone, 0.3% (w/v) yeast extract and 0.01% (w/v) cycloheximide in water, and pH of the medium was adjusted to pH 3.5 with hydrochloric acid (HCl) (Yukphan *et al.*, 2011). Enrichment medium B was composed of 2.0% (w/v) D-glucose, 0.5% (v/v) ethanol, 0.3% (v/v) acetic acid, 1.5% (w/v) peptone, 0.8% (w/v) yeast extract, 0.01% (w/v) cycloheximide in water, and adjusted to pH 3.5 with HCl (Yamada *et al.*, 1999).

Enrichment media observed with microbial growth were inoculated onto solid agar medium containing 2.0% (w/v) D-glucose, 1.0% (v/v) glycerol, 0.5% (v/v) ethanol, 0.5% (w/v) peptone, 0.8% (w/v) yeast extract, 0.7% (w/v) calcium carbonate and 1.5% (w/v) agar in water (Yamada *et al.*, 1999). Inoculated agar plates were incubated aerobically at 28 °C for 7 days. Bacterial colonies of distinctive morphologies on the agar were selected and pure cultures were obtained. All bacterial isolates were routinely maintained on LMG medium 404 (5% (w/v) D-glucose; 1% (w/v) yeast extract and 1.5% (w/v) agar) and frozen glycerol stocks (in 20% (v/v) glycerol) were maintained in −80 °C.

All bacterial isolates were first dereplicated by matrix-assisted laser desorption ionization time-of-flight (MALDI-TOF) mass spectrometry (MS) analysis. Bacterial isolates cultured on LMG medium 404 for 2 days at 28 °C were subjected to ethanol-formic acid protein extraction according to MALDI Biotyper protocol. A saturated α-cyano-4-hydroxycinnamic acid (HCCA) in 50% acetonitrile and 2.5% trifluoroacetic acid was used as matrix solution. Sample prepared on MALDI target plate was analyzed in a Microflex MALDI-TOF bench-top mass spectrometer (Bruker Daltonik GmbH, Leipzig, Germany) using FlexControl software (v3.3). The main spectrum profiles (MSPs) generated were analyzed using MALDI Biotyper Real Time Classification (v3.1) and a cluster analysis based on similarity scorings of these MSPs was done using Biotyper MSP creation method (Goh *et al.*, 2016). From the cluster analysis of MALDI-TOF, representative bacterial isolates from each cluster were selected for preliminary 16S rRNA gene identification.

Subsequently, bacterial isolates in MALDI-TOF cluster 2 that were identified as *O. smaragdina*-associated *Acetobacteraceae* taxon were further dereplicated by random amplified polymorphic DNA (RAPD) fingerprinting. Genomic DNA (gDNA) of bacterial isolate was extracted using MasterPure DNA Purification Kit (EpiCenter, CA, USA) and eluted in Elution Buffer. The RAPD-PCR mixture contained 100 ng of gDNA, 0.45 μM of either primer 270 or primer 272, 200 μM of each dNTPs and 2.5 U of OneTaq™ DNA Polymerase (New England Biolabs) in 25 μL reaction volume. RAPD-PCR was carried out in a Veriti Thermal Cycler (Applied Biosystems) using the following conditions: (i) 1 cycle of 15 min at 95 °C; (ii) 4 cycles of 5 min at 94 °C, 5 min at 36 °C, and 5 min at 72 °C; and (iii) 30 cycles of 1 min at 94 °C, 1 min at 36 °C, and 1 min at 72 °C, followed by (iv) a final extension step at 72 °C for 10 min (Krzewinski *et al.*, 2001). RAPD amplicons were separated by 1% w/v agarose gel electrophoresis. RAPD fingerprints generated were compared in GelJ (Heras *et al.*, 2015) and a cluster analysis using Unweighted Pair Group Method with Arithmetic mean (UPGMA) method was performed with similarity matrix calculated with the Pearson correlation. Bacterial strains Ha5^T^, Ta1 and Jb2 were selected from bacterial isolates of each cluster based on their distinctive RAPD fingerprints.

Reference strains of closely related genera in family *Acetobacteraceae* were included in most analyses. The type strains *Bombella intestini* DSM 28636^T^ and *Saccharibacter floricola* DSM 15669^T^ were acquired from Leibniz Institute-Deutsche Sammlung von Mikroorganismen und Zellkulturen (DSMZ) culture collection. *Neokomagataea thailandica* NBRC 106555^T^, *N. tanensis* NBRC 106556^T^ and *Swingsia samuiensis* NBRC 107927^T^ were acquired from NITE Biological Resource Center (NBRC). Identity of all type strains was confirmed by 16S rRNA gene analysis. All reference strains were routinely maintained on LMG medium 404 and frozen glycerol stocks were maintained in −80 °C.

### Phylogenetic analysis

16S rRNA genes of *Acetobacteraceae* strains were amplified with primers 27F (5’-AGAGTTTGGATCMTGGCTCAG – 3’) and 1492R (5’ – CGGTTACCTTGTTACGACTT – 3’) (Lane, 1991). Amplicons were sent for commercial Sanger sequencing (Apical Scientific, Selangor, Malaysia). The assembled sequences were analyzed by NCBI BLAST tool and 16S-based ID service in EzBioCloud server (http://www.ezbiocloud.net/) (Yoon *et al.*, 2017) for identification. Alignment of 16S rRNA gene sequences with phylogenetically closest genera and species were performed using ClustalW. Phylogenetic trees were constructed in MEGA7 software (Kumar *et al.*, 2016) by neighbourjoining (NJ) and maximum-likelihood (ML) methods. The robustness of tree topologies was estimated by bootstrap analysis (1000 replicates).

### Genomic analysis and calculations of overall genome-relatedness indices

The complete genome of strain Ha5^T^ and draft genomes of strains Ta1 and Jb2 were sequenced using single-molecule real-time sequencing platform (SMRT) (Pacific Biosciences) and Illumina sequencing platform, respectively. Their DNA G + C contents were determined from their whole genome sequences. The NCBI Prokaryotic Genome Annotation Pipeline (PGAP) (Tatusova *et al.*, 2016) and Rapid Annotation using Subsystem Technology (RAST) (Overbeek *et al.*, 2014) were used for genome annotation.

The whole genome sequences of current type members of family *Acetobacteraceae* were obtained from GenBank for further analyses (Supplementary data Table S2). The bacterial identity of these genomes was checked by their 16S rRNA gene sequence using 16S-based ID service in EzBioCloud server (Yoon *et al.*, 2017) and genome completeness was checked with BUSCO v3.0.2 (Simão *et al.*, 2015). Genomes with ambiguous 16S rRNA gene identity and BUSCO genome completeness lesser than 90% were excluded from analysis. Average nucleotide identity (ANI) values between genomes were determined using Orthologous Average nucleotide identity Tool (OAT) (Lee *et al.*, 2016). *In silico* DNA-DNA hybridization (DDH) was calculated using Genome-to-Genome Distance Calculator (GGDC 2.0) (http://ggdc.dsmz.de/distcalc2.php) with the BLAST+ method (Meier-Kolthoff *et al.*, 2013) and the results based on formula 2 were obtained. Average amino acid identity (AAI) between the genomes was determined using CompareM (https://github.com/dparks1134/CompareM) (Konstantinidis and Tiedje, 2005). Percentage of conserved protein (POCP) between the genomes was calculated according to formula by Qin *et al.* (2014).

To elucidate the evolutionary relationships between strains Ha5^T^, Ta1, Jb2 and current taxa of family *Acetobacteraceae*, a phylogenomic analysis was performed as described by Madhaiyan *et al.* (2020). Briefly, panX was used for a core-genome analysis (Ding *et al.*, 2018) in which the groups of homologous genes were identified from all the genomes by similarity search using DIAMOND and rRNA genes were compared by BLASTn, before creating clusters of putatively orthologous genes with Markov Clustering Algorithm (MCL). The core genes were aligned using MAFFT 1.3.7 (Katoh and Standley, 2013) and spurious sequences or poorly aligned regions were removed using trimAl v1.2 (Capella-Gutiérrez *et al.*, 2009). The best-fit substitution model for each nucleotide sequence of the core-genes alignment was determined using ModelTest-NG (Darriba *et al.*, 2020). The core-genome phylogeny was constructed using RAxML-NG, and the robustness of branching was estimated with 100 bootstrap replicates (Kozlov *et al.*, 2019).

### Morphological physiological and biochemical characterizations

Gram staining of bacterial strains was performed using a Gram staining kit (Merck). Strain Ha5^T^ cultured in LMG medium 404 broth for 24 hours was determined for cellular morphology using scanning electron microscope (FEI Quanta FEG 650). Catalase and oxidase activities were assayed using 3% (v/v) hydrogen peroxide and 1% (w/v) *N*, *N*, *N’, N*’-Tetramethyl-1, 4-phenylenediamine (bioMérieux), respectively. Motility was determined on 0.4% semi-solid LMG medium 404 slant (Li *et al.*, 2015, Tittsler and Sandholzer, 1936). The strains were tested for temperature range (4, 10, 15, 20, 25, 30, 35, 37 and 40 °C) for growth on LMG 404 agar medium for 14 days. Tolerance to sodium chloride (NaCl) was determined by growing the strains in LMG medium 404 containing different concentration of NaCl (0, 1, 1.5, 2 and 2.5% (w/v)) (Yun *et al.*, 2017).

Oxidations of lactate and acetate were tested in broth media containing 0.2% (w/v) yeast extract, 0.3% (w/v) peptone, either 0.2% (w/v) of sodium acetate or calcium lactate and 0.002% (w/v) bromothymol blue (Asai *et al.*, 1964). The production of 2-keto-D-gluconic acid and 5-keto-D-gluconic acid from D-glucose was detected using thin layer chromatography as described by Gosselé *et al.* (1980). Production of water-soluble brown pigment was examined on GYC agar containing 2% (w/v) glucose, 2% (w/v) yeast extract, 2% (w/v) calcium carbonate and 2% (w/v) agar (Shimwell *et al.*, 1960). The production of acetic acid from ethanol was tested on agar medium containing 1% (w/v) yeast extract, 2% (w/v) calcium carbonate, 1% (v/v) ethanol and 1.5% (w/v) agar (Shimwell *et al.*, 1960). Bacterial strains were cultured in broth with 1% (w/v) yeast extract and 3% (v/v) glycerol for 7 days at 28 °C and production of dihydroxyacetone from glycerol was tested by addition of Benedict’s solution (Aydin and Aksoy, 2009). Ability of the bacterial strains to utilize ammonium as the sole nitrogen source was tested using Frateur’s modified Hoyer medium containing either 3% (w/v) D-glucose, D-mannitol or 3% (v/v) ethanol (De Ley *et al.*, 1984).

Other biochemical tests including growth of strains in the presence of 30% (w/v) glucose and 1% (w/v) glucose (with modified LMG medium 404 broth), growth in the presence of 0.35% (v/v) acetic acid and in the presence of 1% (w/v) potassium nitrate (KNO_3_) (both in LMG medium 404), growth on 1% (w/v) glutamate agar (Jojima *et al.*, 2004) and growth on mannitol agar (Asai *et al.*, 1964) were tested. Acids production from L-arabinose, D-arabinose, D-glucose, D-galactose, D-mannose, D-fructose, melibiose, sucrose, raffinose, maltose, methanol, D-mannitol, D-sorbitol, glycerol and ethanol were determined in broth medium containing 0.5% (w/v) yeast extract, 0.002% (w/v) bromocresol purple and 1% of the carbon sources (Asai *et al.*, 1964).

### Chemotaxonomic characterizations

Respiratory quinones were analyzed by the Identification Service of the DSMZ, Braunschweig, Germany using a two-stage method (Tindall, 1990a, b). The whole-cell fatty acid methyl esters (FAME) composition was determined using an Agilent Technologies 6890N gas chromatograph (GC). The strains were cultivated on LMG medium 404 agar under aerobic conditions at 28 °C for 48 h following the exact conditions applied on family *Acetobacteraceae* as described by Li *et al.* (2015). Cultures were extracted for fatty acids according to the standard protocol of the Microbial Identification System (Sherlock v6.1; MIDI) (Sasser, 1990). The peaks profiles obtained were identified using the TSBA50 identification library v5.0 (MIDI).

### Nucleotide sequence accession numbers

The 16S rRNA gene sequences of strains Ha5^T^ (accession number MG757796), Ta1 (MN540264) and Jb2 (MN540265) were submitted to GenBank (http://www.ncbi.nlm.nih.gov). The whole genome sequences of strains Ha5^T^, Ta1 and Jb2 were deposited in GenBank under the accession numbers CP038143, SORY00000000 and SORZ00000000.

## Results and Discussion

### Bacterial isolation, dereplication and identification of *Acetobacteraceae* strains

All bacterial isolates obtained were listed in Supplementary data Table S3. Cluster analysis based on MALDI-TOF MS-generated MSPs placed the bacterial isolates into two distantly related clusters in dendrogram created with MALDI Biotyper MSP (Supplementary data Fig. S1). The bacterial isolates in MALDI-TOF MS cluster 1 were placed on the same clade with *Lactobacillus casei* DSM 20011^T^, while bacterial isolates of cluster 2 were placed in same clade with *Gluconobacter oxydans* subspecies *oxydans* B540 UFL in Bruker database. The MALDI-TOF MSPs of bacterial isolates in the same cluster exhibited high similarity and close distance level in MSP dendrogram (Supplementary data Fig. S1).

Representative isolates in MALDI-TOF MS cluster 1 shared highest 16S rRNA gene sequence similarity with *Lactobacillus sanfranciscensis* ATCC 27651^T^ (16S rRNA gene accession number: X76327) in EzBioCloud server (Supplementary data Table S4). On the other hand, representative isolates in MALDI-TOF MS cluster 2 were revealed as one of the *O. smaragdina*-associated *Acetobacteraceae* taxa as they shared highest 16S rRNA gene sequence similarity with *N. tanensis* NBRC 106556^T^ (AB513363) in EzBioCloud server (Supplementary data Table S4).

Bacterial isolates in MALDI-TOF cluster 2 that were obtained from the same *O. smaragdina* sample exhibited highly similar RAPD DNA fingerprints. Based on the cluster analysis using UPGMA method, bacterial isolates from same ant sample were clustered in the same clade, suggesting that these isolates represented a single strain (Supplementary data Fig. S2). Different DNA fingerprints were obtained for bacterial isolates from different ant samples and formed separated clades in RAPD analysis. Strain Jb2 was the only bacterial isolate of the ant sample that belonged to MALDI-TOF cluster 2 which formed individual clade in cluster analysis. Bacterial strains Ha5^T^, Ta1 and Jb2 were selected as the representative strain from each cluster for further analysis.

### Phylogenetic and phylogenomic analyses

Complete 16S rRNA gene sequences were extracted from all available whole genome sequences. The 16S rRNA gene sequence of strain Ha5^T^ shared 99.93% similarity with strains Ta1 and Jb2 and was only different by 1 bp (Supplementary data Table S5). The strains shared highest 16S rRNA gene sequence similarities with two members of the genus *Neokomagataea*, *N. tanensis* NBRC 106556^T^ (94.56-94.63%) and *N. thailandica* NBRC 106555^T^ (94.03-94.10%) and *B. intestini* DSM 28636^T^ (94.09-94.16%) (Supplementary data Table S5). However, strains Ha5^T^, Ta1 and Jb2 formed a cluster phylogenetically apart from genus *Neokomagataea* in phylogenetic trees calculated with ML and NJ methods. Instead, both phylogenetic trees grouped the bacterial strains in a distinct clade which was closer to a separated clade that included *S’. floricola* DSM 15669^T^ and two members of *Bombella,* before placing the mentioned taxa in a larger clade that included genera *Neokomagataea* and *Swingsia samuiensis* NBRC 107927^T^ (Supplementary data Fig. S3 and S4). In short, strains Ha5^T^, Ta1 and Jb2 formed a distinct lineage based on 16S rRNA gene phylogenetic analysis of family *Acetobacteraceae* which suggested that they should be differentiated as a new genus.

Although 16S rRNA gene remains the widely used molecular marker in taxonomic and phylogenetic study, the approach omits the overall genetic differences especially among closely related species. To obtain a robust construction of phylogeny, we therefore decided to elucidate the phylogenetic relationship of strains Ha5^T^, Ta1, Jb2 and family *Acetobacteraceae* members through phylogenomic analysis. A total of 75 whole genome sequences of strains were acquired from this study or from GenBank (Supplementary data Table S2) was included for phylogenomic analysis using panX. Genomes that were without 16S rRNA gene, showed ambiguous 16S rRNA gene identity and BUSCO genome completeness lower than 90% were excluded from analysis.

The phylogenomic analysis revealed the existence of 96 orthologous gene clusters which were identified as the core genes from all genomes included in this study. The ML phylogenetic tree based on these concatenated genes (total 76641 base pairs) showed placement of all genera in family *Acetobacteraceae* on distinctive phylogenetic lineages (Fig. 1). Similar to phylogenetic trees based on 16S rRNA gene sequences, strains Ha5^T^, Ta1 and Jb2 formed a distinct cluster in the phylogenomic tree and were placed in the same clade with *S. floricola* DSM 15669^T^ and *B. intestini* DSM 28636^T^. These analyses showed that all nodes were supported by bootstrap values greater than 50 which indicated confident robustness of the result. The phylogenomic tree corroborated the phylogenetic distinction of strains Ha5^T^, Ta1 and Jb2 from all members of the family *Acetobacteraceae* and indicated that they should be classified into a novel genus in the family.

**Fig. 1.**
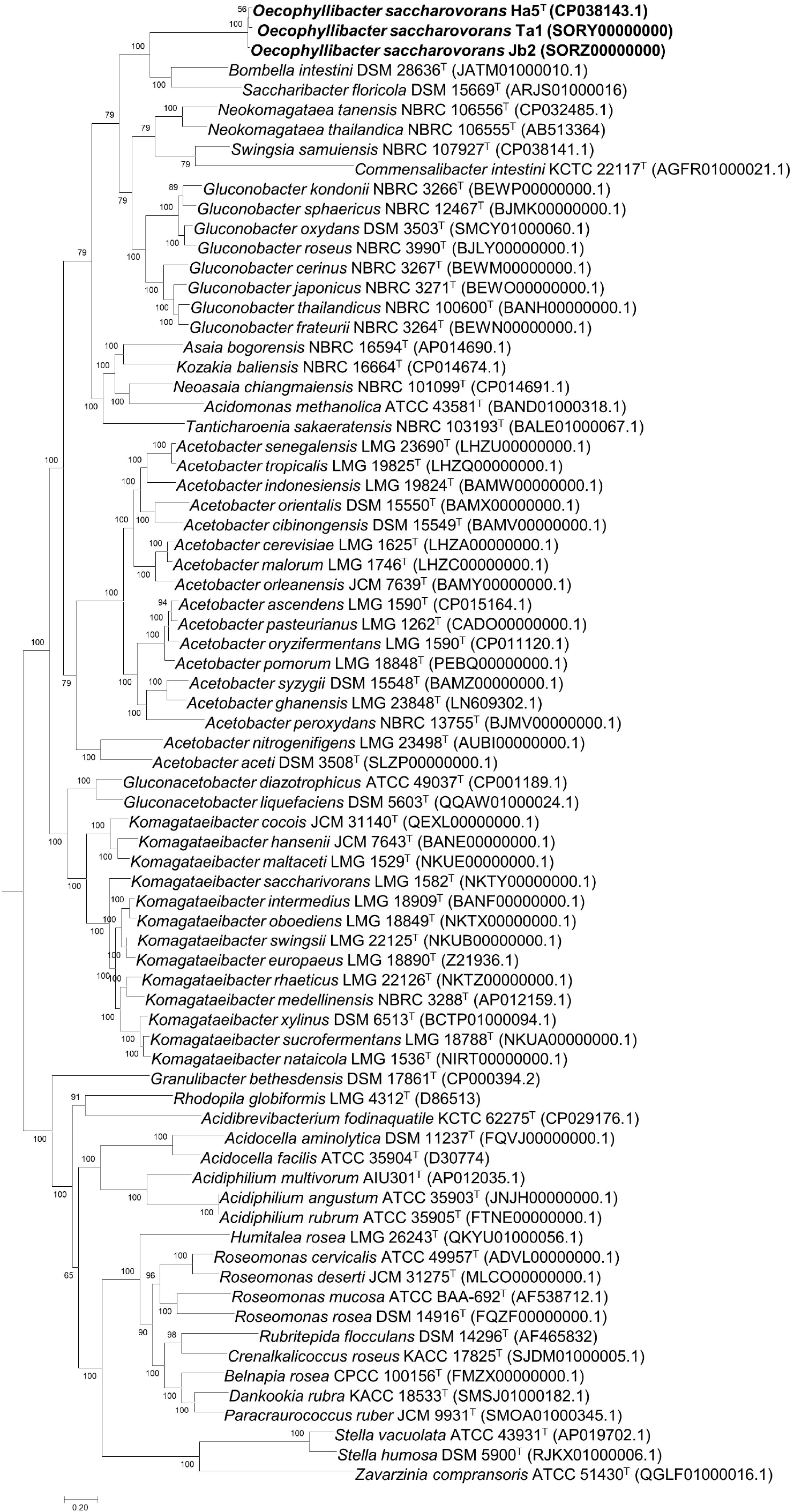
Phylogenomic tree of 96 concatenated core gene sequences revealed from whole genomes of family *Acetobacteraceae* showing the relationship of strains Ha5^T^, Ta1, Jb2 and their closest relatives in the family *Acetobacteraceae*. The tree was constructed using maximum likelihood method. The values at the nodes indicate bootstrap values expressed as percentages of 100 replications while the bar length indicates 0.20 substitutions per site.

### Genomic analysis

The whole genomes of strains Ha5^T^, Ta1 and Jb2 were 1.92 Mb to 1.95 Mb in size and exhibited proximate genomic DNA G + C contents in the range of 61.56 to 61.65 mol% (Supplementary data Table S2). The list of genomes for current *Acetobacteraceae* taxa obtained in this study or from GenBank showed large range in terms of size (1.92 Mb to 7.78 Mb), DNA G + C content (36.8 mol% to 73.9 mol%), number of genes (1,632 to 7,451) and number of protein coding genes (1,574 to 7,102) (Supplementary data Table S2). Strains Ha5^T^, Ta1 and Jb2 shared pairwise ANI values in the range of 97.5-97.7% (Supplementary data Fig. S5), pairwise *in silico* DDH values from 77.4% to 79.3% (Supplementary data Fig. S6), and AAI values in the range of 97.9 to 98.1% (Supplementary data Fig. S7). These values are higher than the cut-off values (95–96% for ANI; >70% for *in silico* DDH; >95% for AAI) proposed for species boundary and thus showed that these strains belonged to the same species (Konstantinidis and Tiedje, 2005, Richter and Rosselló-Móra, 2009).

Furthermore, the ANI, *in silico* DDH and AAI values between the strains Ha5^T^, Ta1 and Jb2 and other members of family *Acetobacteraceae* were far below the proposed cut-off values for species delineation (≤70.7% for ANI; ≤37.1% for *in silico* DDH; ≤66.0% for AAI) (Supplementary data Fig. S5, S6, S7). It is also noteworthy that the ≤66.0% pairwise AAI values between strains Ha5^T^, Ta1, Jb2 and type members of family *Acetobacteraceae* were within the reported range of genus-level pairwise AAI difference (65% to 72% AAI values) (Konstantinidis and Tiedje, 2005). Through these indices, it was substantiated that strains Ha5^T^, Ta1, Jb2 did not belong to any of the current species in family *Acetobacteraceae* and should be delineated as a single species within a distinct genus in the family. Overall, these findings further supported that the strains Ha5^T^, Ta1 and Jb2 represent a novel genus of the family *Acetobacteraceae*.

On the other hand, we had looked into POCP that was suggested as an alternative prokaryotic genus delineation index (Qin *et al.*, 2014). Based on the proposed formula, strains Ha5^T^, Ta1 and Jb2 shared high POCP values in the range of 97.2–97.6% (Supplementary data Fig. S8). However, the bacterial strains also shared higher than 50% POCP values with many type members of family *Acetobacteraceae,* including *Bombella, Neokomagataea, Swingsia, Saccharibacter, Gluconobacter, Acetobacter* and *Asaia* (Supplementary data Fig. S8). The high POCP values could be attributed to large difference in number of protein coding genes between strains Ha5^T^, Ta1 and Jb2 and species of other genera. With reference to the suggested cut-off threshold, two bacterial strains with a POCP value of higher than 50% between them can be defined as species grouped in a prokaryotic genus (Qin *et al.*, 2014). The result contradicted with calculations of ANI, *in silico* DDH and AAI which revealed that strains Ha5^T^, Ta1 and Jb2 did not belong to any of the species of existing genera of family *Acetobacteraceae*. Besides, the suggested cut-off threshold also did not apply to many other members between existing genera of family *Acetobacteraceae* (Supplementary data Fig. S8). Our findings showed that POCP had disputable inter-genera boundary and may not be reliable for genus-level delineation of family *Acetobacteraceae*.

Strains Ha5^T^, Ta1 and Jb2 showed highly similar gene distribution into different RAST subsystem categories. The categories for protein metabolism, amino acids and derivatives and cofactors, vitamins, prosthetic groups and pigments were assigned with highest number of genes (in descending order) (Supplementary data Fig. S9). The genome annotation detected the presence of a catalase gene in their genomes which confirmed their catalase-positive activity (Supplementary data Table S6). Besides, genes encoding for cytochrome *d* ubiquinol oxidase subunit I and II were predicted but not gene encoding for cytochrome *c* oxidase (Supplementary data Table S6). This finding corroborated the absence of cytochrome *c* oxidase in the bacterial strains and thus their negative oxidase activity (Table 1). The genomes of strains Ha5^T^, Ta1 and Jb2 revealed the presence of biosynthetic genes for cofactors and vitamins including riboflavin, biotin, folate, pantothenate, riboflavin, thiamine and pyridoxine (Supplementary data Table S6). These genes are crucial in nutritional contribution to the ant host *O. smaragdina*. Among the mentioned vitamins and cofactors, riboflavin is an absolute dietary supplement for higher animals that do not possess the riboflavin biosynthetic pathway. As riboflavin is only synthesized by plants and most bacteria, *O. smaragdina* could be therefore relying on dietary riboflavin or provisioning by strains Ha5^T^, Ta1 and Jb2 through biosynthesis.

**Table 1.**
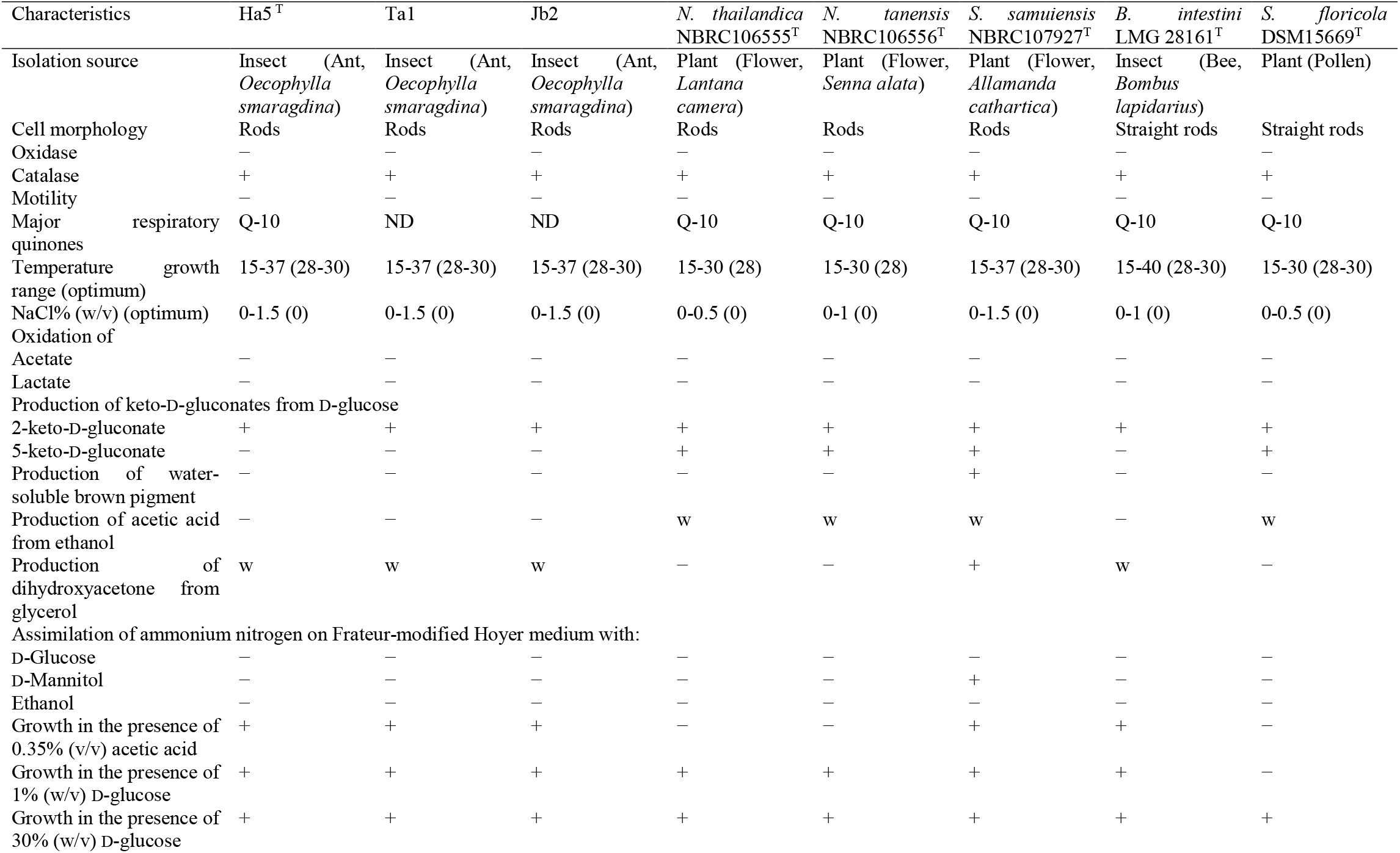
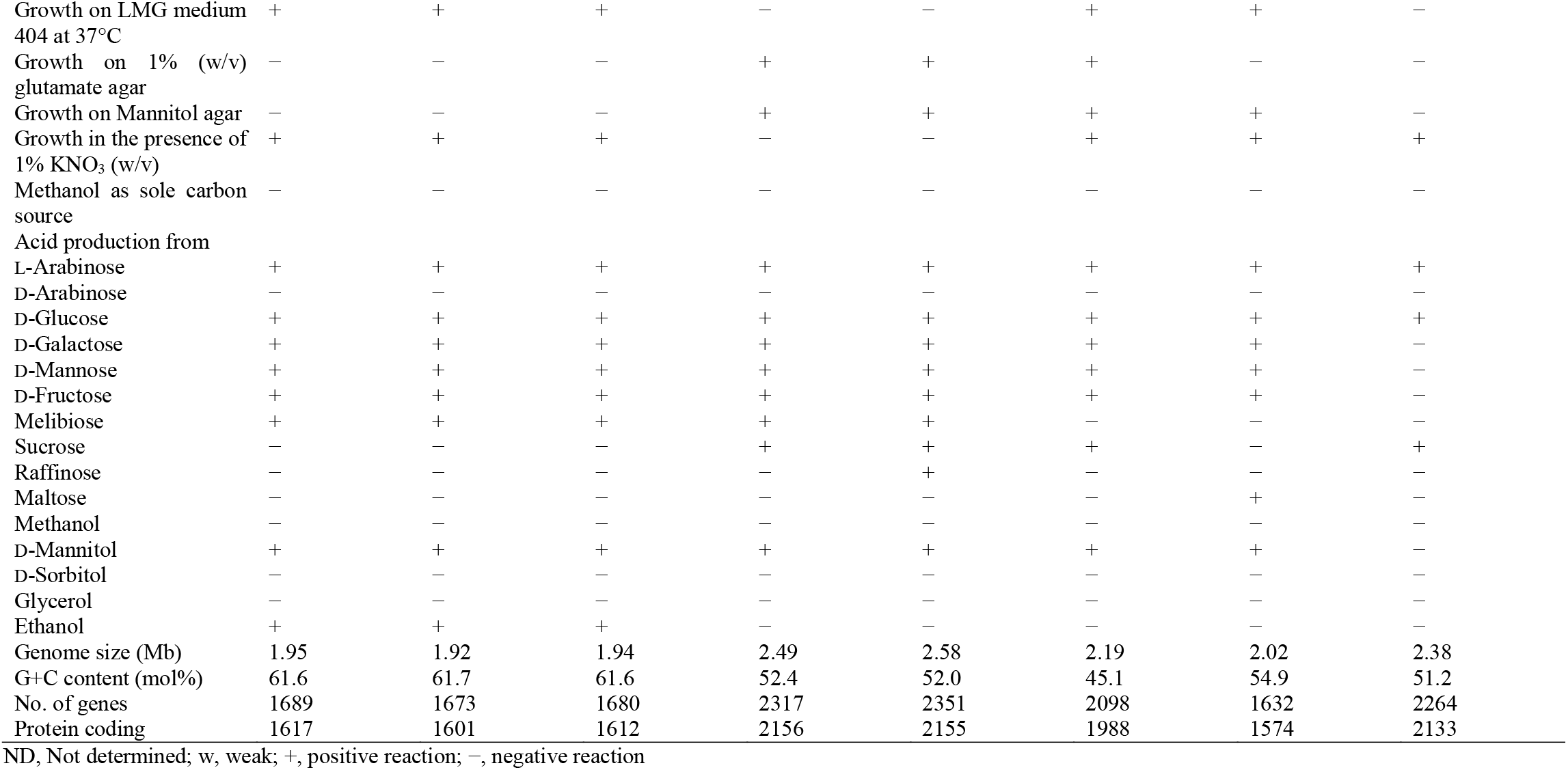
Differential characteristics between bacterial strains Ha5^T^, Ta1, Jb2 and type strains of closely related genera of family *Acetobacteraceae*. All results were obtained in this study.

In addition, genes involved in biosynthesis for all non-essential amino acids and multiple essential amino acids (except for isoleucine, leucine and valine) were identified in the genomes of strains Ha5^T^, Ta1 and Jb2 (Supplementary data Table S6). It is suggested the strains supply amino acids to the ant host that relies mostly on plant sap and honeydew which constitute an unbalanced diet lacking of amino acids (Holldobler and Wilson, 1990). However, the omnivorous *O. smaragdina* also preys on intruding insects which could be the source of nitrogen (Peng and Christian, 2004). The lacking of biosynthetic genes in the strains may imply a less important role in supplying essential amino acids to the host. Nonetheless, bacterial symbionts are recognized for biosynthesis and supply of amino acids to their insect hosts. Notably, the endosymbiont *Blochmannia* sp. in ants of the genus *Camponotus* retains biosynthetic genes for amino acids including tyrosine even with a strongly reduced genome (Feldhaar *et al.*, 2007). Overall, the genomic analysis provided insights into biosynthetic genes of strains Ha5^T^, Ta1 and Jb2 and revealed their biological functions as symbiont of *O. smaragdina.*

### Morphological, physiological and biochemical characteristics

Strains Ha5^T^, Ta1 and Jb2 formed beige-colored, shiny, circular, convex, smooth and opaque small colonies on LMG medium 404 agar after incubation for 2 days at 28 °C. The cells were Gram-stain negative and rod-shaped when observed under the light microscope. From SEM micrograph, the rod-shaped cells of strain Ha5^T^ had a size of 0.45-0.5 μm in width and 0.8-1.0 μm in length (Supplementary data Fig. S10). No flagella were observed and all strains were non-motile on semi-solid agar.

Phenotypic and physiological tests performed on strains Ha5^T^, Ta1 and Jb2 showed similar results and their physiological differentiation with all reference strains are listed in Table 1. Similar to the closest neighbors in family *Acetobacteraceae*, strains Ha5^T^, Ta1 and Jb2 were tested negative for oxidase activity but positive for catalase activity. They did not carry out oxidation of lactate and acetate. Strains Ha5^T^, Ta1 and Jb2 showed glucose requirement for growth and were able to grow in media with 1% and 30% (w/v) glucose (Table 1). While growth was observed in temperature from 15 to 37 °C, optimum growth occurred aerobically at 28-30 °C. No growth occurred at 10 °C and 40 °C. All three strains showed growth in medium containing NaCl up to 1.5% (w/v) but optimum growth occurred at 0%. Phenotypic features that differentiate strain Ha5^T^ and closest neighbors of family *Acetobacteraceae* included the production of 5-keto-D-gluconate from D-glucose, production of water soluble brown pigment, production of dihydroxyacetone from glycerol, growth in the presence of 0.35% v/v acetic acid, growth in the presence of 1% KNO_3_ (w/v) and acid production from various carbon sources (Table 1).

### Chemotaxonomic characteristics

Analysis of respiratory quinones revealed ubiquinone-10 (Q-10) as the major ubiquinone in strain Ha5^T^ (Table 1). This is consistent with all members of family *Acetobacteraceae* except genus *Acetobacter* that is having ubiquinone-9 (Q-9) as the major ubiquinone (Yamada *et al.*, 1968, Yamada and Yukphan, 2008). The major cellular fatty acid profile of the Ha5^T^ contained C_18:1_ *ω7c* (36.11%), C_16:0_ (22.31%) and C_19:0_ *ω8c* cyclo (18.43%) which accounted for 76.85% of the total fatty acids (Table 2). Strain Ha5^T^ has similar overall major fatty acid profile to those of strains Ta1 and Jb2 except differences in respective proportions of some fatty acid components in low abundance (Table 2). Strain Ha5^T^ and all reference strains have C_18:1_ *ω7c* and C_16:0_ as the major fatty acids and differences are observed mostly with the low abundance fatty acids (Table 2).

**Table 2.**
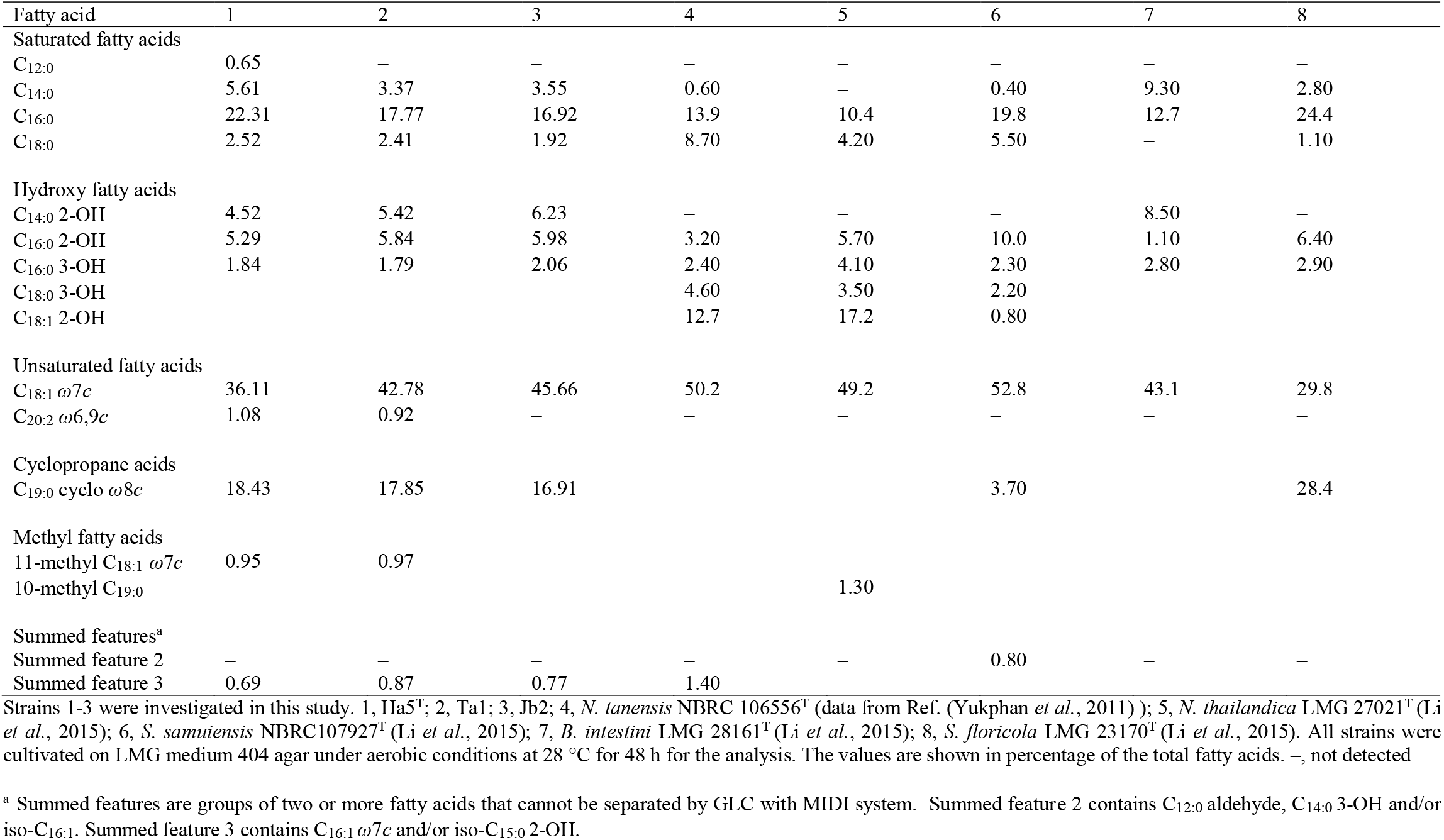
Fatty acid composition of strains Ha5^T^, Ta1, Jb2 and type strains of closely related genera in family *Acetobacteraceae*.

| Fatty acid | 1 | 2 | 3 | 4 | 5 | 6 | 7 | 8 |
| --- | --- | --- | --- | --- | --- | --- | --- | --- |
| Saturated fatty acids |  |  |  |  |  |  |  |  |
| C<sub>12:0</sub> | 0.65 | — | — | — | — | — | — | — |
| C<sub>14:0</sub> | 5.61 | 3.37 | 3.55 | 0.60 | — | 0.40 | 9.30 | 2.80 |
| C<sub>16:0</sub> | 22.31 | 17.77 | 16.92 | 13.9 | 10.4 | 19.8 | 12.7 | 24.4 |
| C<sub>18:0</sub> | 2.52 | 2.41 | 1.92 | 8.70 | 4.20 | 5.50 | — | 1.10 |
| Hydroxy fatty acids |  |  |  |  |  |  |  |  |
| C<sub>14:0</sub> 2-OH | 4.52 | 5.42 | 6.23 | — | — | — | 8.50 | — |
| C<sub>16:0</sub> 2-OH | 5.29 | 5.84 | 5.98 | 3.20 | 5.70 | 10.0 | 1.10 | 6.40 |
| C<sub>16:0</sub> 3-OH | 1.84 | 1.79 | 2.06 | 2.40 | 4.10 | 2.30 | 2.80 | 2.90 |
| C<sub>18:0</sub> 3-OH | — | — | — | 4.60 | 3.50 | 2.20 | — | — |
| C<sub>18:1</sub> 2-OH | — | — | — | 12.7 | 17.2 | 0.80 | — | — |
| Unsaturated fatty acids |  |  |  |  |  |  |  |  |
| C<sub>18:1</sub> ω7c | 36.11 | 42.78 | 45.66 | 50.2 | 49.2 | 52.8 | 43.1 | 29.8 |
| C<sub>20:2</sub> ω6,9c | 1.08 | 0.92 | — | — | — | — | — | — |
| Cyclopropane acids |  |  |  |  |  |  |  |  |
| C<sub>19:0</sub> cyclo ω8c | 18.43 | 17.85 | 16.91 | — | — | 3.70 | — | 28.4 |
| Methyl fatty acids |  |  |  |  |  |  |  |  |
| 11-methyl C<sub>18:1</sub> ω7c | 0.95 | 0.97 | — | — | — | — | — | — |
| 10-methyl C<sub>19:0</sub> | — | — | — | — | 1.30 | — | — | — |
| Summed features<sup>a</sup> |  |  |  |  |  |  |  |  |
| Summed feature 2 | — | — | — | — | — | 0.80 | — | — |
| Summed feature 3 | 0.69 | 0.87 | 0.77 | 1.40 | — | — | — | — |

Strains 1-3 were investigated in this study. 1, Ha5<sup>T</sup>; 2, Ta1; 3, Jb2; 4, *N. tanensis* NBRC 106556<sup>T</sup> (data from Ref. (Yukphan *et al.*, 2011) ); 5, *N. thailandica* LMG 27021<sup>T</sup> (Li *et al.*, 2015); 6, *S. samuiensis* NBRC107927<sup>T</sup> (Li *et al.*, 2015); 7, *B. intestini* LMG 28161<sup>T</sup> (Li *et al.*, 2015); 8, *S. floricola* LMG 23170<sup>T</sup> (Li *et al.*, 2015). All strains were cultivated on LMG medium 404 agar under aerobic conditions at 28 °C for 48 h for the analysis. The values are shown in percentage of the total fatty acids. —, not detected
<sup>a</sup> Summed features are groups of two or more fatty acids that cannot be separated by GLC with MIDI system. Summed feature 2 contains C<sub>12:0</sub> aldehyde, C<sub>14:0</sub> 3-OH and/or iso-C<sub>16:1</sub>. Summed feature 3 contains C<sub>16:1</sub> ω7c and/or iso-C<sub>15:0</sub> 2-OH.

### Taxonomic conclusion

Taxonomic characterizations of bacterial strains Ha5^T^, Ta1 and Jb2 revealed that they were a member of the family *Acetobacteraceae*. These bacterial strains were genomically distinctive based on RAPD DNA fingerprinting. They showed high phenotypic stability based on biochemical, physiological and chemotaxonomic characterizations, high genomic coherence as indicated by calculations of various overall genome-relatedness indices, and were placed in a close monophyletic cluster in 16S rRNA gene-based phylogenetic analysis and core genome-based phylogenomic analysis. These findings showed that bacterial strains Ha5^T^, Ta1 and Jb2 should belong to the same species. Besides, it was suggested in a unified bacterial taxonomic framework that 94.5% 16S rRNA sequence similarity provides strong evidence that two taxa should be distinct at genus level but 96% similarity could be considered as the threshold if it is supported by phenotypic, phylogenetic and other data (Yarza *et al.*, 2014). Bacterial strains Ha5^T^, Ta1 and Jb2 shared highest 16S rRNA gene sequence similarity at ≤94.63% with the closest neighbour *N. tanensis* NBRC 106556^T^. Based on their distinctive phenotypic characteristics, low genomic coherence and distinctive placement in a monophyletic lineage from all current taxa of family *Acetobacteraceae,* strains Ha5^T^, Ta1 and Jb2 should represent a novel species of a novel genus within family *Acetobacteraceae,* for which the name *Oecophyllibacter saccharovorans* gen. nov., sp. nov. is proposed.

### Descriptions of *Oecophyllibacter* gen. nov

*Oecophyllibacter* (Oe.co.phyl.li.bac.ter. N.L. fem. n. *Oecophylla* a weaver ant genus; N.L. masc. n. *bacter* a rod; N.L. masc. n. *Oecophyllibacter,* a rod from the weaver ant *Oecophylla).*

Cells are Gram-negative staining, aerobic, non-motile, rod-shaped and occur singly. Catalase positive and oxidase negative. Major respiratory quinone is ubiquinone Q-10. Predominant cellular fatty acids are C_18:1_ *ω7c,* C_16:0_, C_19:0_ *ω8c* cyclo, C14:0 and C_16:0_ 2-OH. Chemoorganotrophic metabolism. The genomic G + C content is in the range of 61.56–61.65 mol%. The type species is *Oecophyllibacter saccharovorans*.

### Descriptions of *Oecophyllibacter saccharovorans* sp. nov

*Oecophyllibacter saccharovorans* (sac.cha.ro.vo’rans. Gr. fem. n. *sakchar* sugar; L. pres. part. *vorans* eating; N.L. part adj. *saccharovorans* sugar eating).

Displays the following properties in addition to those described for the genus. Cells are rod-shaped, 0.45-0.5 μm in width and 0.8-1.0 μm in length after 24 hr incubation in LMG medium 404 broth. Colonies are beige-colored, small (approximately 1-3 mm), circular, convex, shiny and smooth with entire margin after 2 days incubation at 28 °C on LMG medium 404. Growth occurs at 15-37 °C (optimum 28-30 °C) and with 0-1.5% (w/v) NaCl (optimum 0% (w/v)). Requires glucose for growth and able to grow in 1% and 30% glucose (w/v) media with 1% yeast extract. Able to grow in the presence of 0.35% (v/v) acetic acid and 1% KNO_3_ (w/v). Produces 2-keto-D-gluconic acid but not 5-keto-D-gluconic acid from D-glucose. Does not oxidize ethanol to acetic acid. Does not oxidize lactate and acetate to carbon dioxide and water. No growth with ammonium as the sole nitrogen source. Does not produce water-soluble brown pigment. Weak in production of dihydroxyacetone from glycerol. No growth on 1% glutamate agar. Does not utilize methanol as sole carbon source. Produces acids from L-arabinose, D-glucose, D-galactose, D-mannose, D-fructose, melibiose, D-mannitol and ethanol, but not from D-arabinose, sucrose, raffinose, maltose, methanol, D-sorbitol and glycerol. The genomic G + C content of the type strain is 61.56 mol% based on the complete genome sequence. The type strain Ha5^T^ (= DSM106907^T^ = LMG30589^T^ = NBRC 113643^T^ = KCTC62952^T^) was isolated from weaver ant *O. smaragdina* in Kuala Lumpur, Malaysia.

## Acknowledgements

The authors are grateful to Professor Aharon Oren for his advice on etymology and nomenclature of the strains. KOC thanks MyBrain15 Postgraduate Scholarship Programme for the scholarship (MyPhD, KPT(B)900909146137). WSST thanks the Bright Sparks Program of the University of Malaya for scholarship awarded. This work was supported by University of Malaya Research Grants (FRGS grant FP022-2018A), University of Malaya High Impact Research Grants (UM-MOHE HIR Grant UM.C/625/1/HIR/MOHE/ CHAN/14/1, Grant No. H-50001-A000027; UM-MOHE HIR Grant UM.C/625/1/HIR/MOHE/CHAN/01, Grant No. A-000001-50001) awarded to KGC and Postgraduate Research (PPP) Grant (Grant No. PG089-2015B) awarded to KOC.

## References

Asai, T., Iizuka, H., and Komagata, K. 1964. The flagellation and taxonomy of genera *Gluconobacter* and *Acetobacter* with reference to the existence of intermediate strains. The Journal of General and Applied Microbiology. 10, 95–126.

Aydin, Y. A. and Aksoy, N. D. 2009. Isolation of cellulose producing bacteria from wastes of vinegar fermentation. Presented at the WCECS 2009: World congress on engineering and computer science, Hong Kong.

Brown, B. P. and Wernegreen, J. J. 2019. Genomic erosion and extensive horizontal gene transfer in gut-associated *Acetobacteraceae*. BMC Genomics. 20, 472.

Capella-Gutiérrez, S., Silla-Martínez, J. M., and Gabaldón, T. 2009. trimAl: a tool for automated alignment trimming in large-scale phylogenetic analyses. Bioinformatics. 25, 1972–1973.

Chua, K.-O., Song, S.-L., Yong, H.-S., See-Too, W.-S., Yin, W.-F., and Chan, K.-G. 2018. Microbial Community Composition Reveals Spatial Variation and Distinctive Core Microbiome of the Weaver Ant *Oecophylla smaragdina* in Malaysia. Sci. Rep. 8, 10777.

Cleenwerck, I., Camu, N., Engelbeen, K., De Winter, T., Vandemeulebroecke, K., De Vos, P., and De Vuyst, L. 2007. *Acetobacter ghanensis* sp. nov., a novel acetic acid bacterium isolated from traditional heap fermentations of Ghanaian cocoa beans. Int. J. Syst. Evol. Microbiol. 57, 1647–1652.

Crotti, E., Chouaia, B., Alma, A., Favia, G., Bandi, C., Bourtzis, K., and Daffonchio, D. 2016. Acetic acid bacteria as symbionts of insects, p. 121–142. Acetic Acid Bacteria, Springer.

Crotti, E., Rizzi, A., Chouaia, B., Ricci, I., Favia, G., Alma, A., Sacchi, L., Bourtzis, K., Mandrioli, M., and Cherif, A. 2010. Acetic acid bacteria, newly emerging symbionts of insects. Appl. Environ. Microbiol. 76, 6963–6970.

Crotti, E., Sansonno, L., Prosdocimi, E. M., Vacchini, V., Hamdi, C., Cherif, A., Gonella, E., Marzorati, M., and Balloi, A. 2013. Microbial symbionts of honeybees: A promising tool to improve honeybee health. New Biotechnol. 30, 716–722.

Darriba, D., Posada, D., Kozlov, A. M., Stamatakis, A., Morel, B., and Flouri, T. 2020. ModelTest-NG: a new and scalable tool for the selection of DNA and protein evolutionary models. Mol. Biol. Evol. 37, 291–294.

De Ley, J., Gillis, M., and Swings, J. 1984. Family VI. *Acetobacteraceae*, p. 267–278. *In* N. R. Krieg and J. G. Holt (eds.), Bergey’s manual of systematic bacteriology, The Williams & Wilkins Co., Baltimore, USA.

Ding, W., Baumdicker, F., and Neher, R. A. 2018. panX: pan-genome analysis and exploration. Nucleic Acids Res. 46, e5.

Favia, G., Ricci, I., Damiani, C., Raddadi, N., Crotti, E., Marzorati, M., Rizzi, A., Urso, R., Brusetti, L., and Borin, S. 2007. Bacteria of the genus *Asaia* stably associate with *Anopheles stephensi*, an Asian malarial mosquito vector. Proc. Natl. Acad. Sci. 104, 9047–9051.

Feldhaar, H., Straka, J., Krischke, M., Berthold, K., Stoll, S., Mueller, M. J., and Gross, R. 2007. Nutritional upgrading for omnivorous carpenter ants by the endosymbiont *Blochmannia*. BMC Biol. 5, 48.

Gillis, M. and De Ley, J. 1980. Intra-and intergeneric similarities of the ribosomal ribonucleic acid cistrons of Acetobacter and Gluconobacter. Int. J. Syst. Evol. Microbiol. 30, 7–27.

Goh, S.-Y., Khan, S. A., Tee, K. K., Kasim, N. H. A., Yin, W.-F., and Chan, K.-G. 2016. Quorum sensing activity of Citrobacter amalonaticus L8A, a bacterium isolated from dental plaque. Sci. Rep. 6, 20702.

Gomes, R. J., Borges, M. D. F., Rosa, M. D. F., Castro-Gómez, R. J. H., and Spinosa, W. A. 2018. Acetic acid bacteria in the food industry: Systematics, characteristics and applications. Food Technol. Biotechnol. 56, 139–151.

Gosselé, F., Swings, J., and De Ley, J. 1980. A rapid, simple and simultaneous detection of 2-keto-, 5-keto-and 2, 5-diketogluconic acids by thin-layer chromatography in culture media of acetic acid bacteria. Zentralblatt für Bakteriologie: I. Abt. Originale C: Allgemeine, angewandte und ökologische Mikrobiologie. 1, 178–181.

Heras, J., Domínguez, C., Mata, E., Pascual, V., Lozano, C., Torres, C., and Zarazaga, M. 2015. GelJ-a tool for analyzing DNA fingerprint gel images. BMC Bioinformatics. 16, 270.

Holldobler, B. and Wilson, E. O. 1990. The ants, p. Pages. Harvard University Press, Massachusetts, United States.

Jojima, Y., Mihara, Y., Suzuki, S., Yokozeki, K., Yamanaka, S., and Fudou, R. 2004. *Saccharibacter floricola* gen. nov., sp. nov., a novel osmophilic acetic acid bacterium isolated from pollen. Int. J. Syst. Evol. Microbiol. 54, 2263–2267.

Katoh, K. and Standley, D. M. 2013. MAFFT multiple sequence alignment software version 7: Improvements in performance and usability. Mol. Biol. Evol. 30, 772–780.

Kersters, K., Lisdiyanti, P., Komagata, K., and Swings, J. 2006. The family *Acetobacteraceae*: the genera *Acetobacter, Acidomonas, Asaia, Gluconacetobacter, Gluconobacter*, and *Kozakia*, p. 163–200. *In* M. Dworkin, S. Falkow, E. Rosenberg, K. H. Schleifer and E. Stackebrandt (eds.), The Prokaryotes, Springer, New York, NY.

Kim, K. H., Cho, G. Y., Chun, B. H., Weckx, S., Moon, J. Y., Yeo, S.-H., and Jeon, C. O. 2018. *Acetobacter oryzifermentans* sp. nov., isolated from Korean traditional vinegar and reclassification of the type strains of *Acetobacter pasteurianus* subsp. *ascendens* (Henneberg 1898) and *Acetobacter pasteurianus* subsp. *paradoxus* (Frateur 1950) as *Acetobacter ascendens* sp. nov., comb. nov. Syst. Appl. Microbiol. 41, 324–332.

Konstantinidis, K. T. and Tiedje, J. M. 2005. Towards a genome-based taxonomy for prokaryotes. J. Bacteriol. 187, 6258–6264.

Kounatidis, I., Crotti, E., Sapountzis, P., Sacchi, L., Rizzi, A., Chouaia, B., Bandi, C., Alma, A., Daffonchio, D., and Mavragani-Tsipidou, P. 2009. *Acetobacter tropicalis* is a major symbiont of the olive fruit fly *(Bactrocera oleae)*. Appl. Environ. Microbiol. 75, 3281–3288.

Kozlov, A. M., Darriba, D., Flouri, T., Morel, B., and Stamatakis, A. 2019. RAxML-NG: a fast, scalable and user-friendly tool for maximum likelihood phylogenetic inference. Bioinformatics. 35, 4453–4455.

Krzewinski, J. W., Nguyen, C. D., Foster, J. M., and Burns, J. L. 2001. Use of random amplified polymorphic DNA PCR to examine epidemiology of *Stenotrophomonas maltophilia* and *Achromobacter (Alcaligenes) xylosoxidans* from patients with cystic fibrosis. J. Clin. Microbiol. 39, 3597–3602.

Kumar, S., Stecher, G., and Tamura, K. 2016. MEGA7: Molecular Evolutionary Genetics Analysis version 7.0 for bigger datasets. Mol. Biol. Evol. 33, 1870–1874.

Lane, D. J. 1991. 16S/23S rRNA sequencing, p. 115–175. *In* E. S. a. M. Goodfellow (ed.), Nucleic acid techniques in bacterial systematics, Wiley & Sons, Chichester, United Kingdom.

Lee, I., Kim, Y. O., Park, S.-C., and Chun, J. 2016. OrthoANI: An improved algorithm and software for calculating average nucleotide identity. Int. J. Syst. Evol. Microbiol. 66, 1100–1103.

Li, L., Praet, J., Borremans, W., Nunes, O. C., Manaia, C. M., Cleenwerck, I., Meeus, I., Smagghe, G., De Vuyst, L., and Vandamme, P. 2015. *Bombella intestini* gen. nov., sp. nov., an acetic acid bacterium isolated from bumble bee crop. Int. J. Syst. Evol. Microbiol. 65, 267–273.

Lin, J. Y., Russell, J. A., Sanders, J. G., and Wertz, J. T. 2016. *Cephaloticoccus* gen. nov., a new genus of ‘Verrucomicrobia’containing two novel species isolated from *Cephalotes* ant guts. Int. J. Syst. Evol. Microbiol. 66, 3034–3040.

Madhaiyan, M., Saravanan, V. S., and See-Too, W.-S. 2020. Genome-based analyses reveal the presence of 12 heterotypic synonyms in the genus Streptomyces and emended descriptions of Streptomyces bottropensis, Streptomyces celluloflavus, Streptomyces fulvissimus, Streptomyces glaucescens, Streptomyces murinus, and Streptomyces variegatus. Int. J. Syst. Evol. Microbiol. ijsem004217.

Meier-Kolthoff, J. P., Auch, A. F., Klenk, H.-P., and Göker, M. 2013. Genome sequence-based species delimitation with confidence intervals and improved distance functions. BMC Bioinformatics. 14, 60.

Overbeek, R., Olson, R., Pusch, G. D., Olsen, G. J., Davis, J. J., Disz, T., Edwards, R. A., Gerdes, S., Parrello, B., and Shukla, M. 2014. The SEED and the Rapid Annotation of microbial genomes using Subsystems Technology (RAST). Nucleic Acids Res. 42, D206–D214.

Peng, R. K. and Christian, K. 2004. The weaver ant, *Oecophylla smaragdina* (Hymenoptera: Formicidae), an effective biological control agent of the red-banded thrips, *Selenothrips rubrocinctus* (Thysanoptera: Thripidae) in mango crops in the Northern Territory of Australia. Int. J. Pest Manage. 50, 107–114.

Peng, R. K., Christian, K., and Gibb, K. 1995. The effect of the green ant, *Oecophylla smaragdina* (Hymenoptera: Formicidae), on insect pests of cashew trees in Australia. Bull. Entomol. Res. 85, 279–284.

Pimid, M., Hassan, A., Tahir, N. A., and Thevan, K. 2014. Colony Structure of the Weaver Ant, *Oecophylla smaragdina* (Fabricius)(Hymenoptera: Formicidae). Sociobiol. 59, 1–10.

Qin, Q.-L., Xie, B.-B., Zhang, X.-Y., Chen, X.-L., Zhou, B.-C., Zhou, J., Oren, A., and Zhang, Y.-Z. 2014. A proposed genus boundary for the prokaryotes based on genomic insights. J. Bacteriol. 196, 2210–2215.

Richter, M. and Rosselló-Móra, R. 2009. Shifting the genomic gold standard for the prokaryotic species definition. Proc. Natl. Acad. Sci. 106, 19126–19131.

Roh, S. W., Nam, Y.-D., Chang, H.-W., Kim, K.-H., Kim, M.-S., Ryu, J.-H., Kim, S.-H., Lee, W.-J., and Bae, J.-W. 2008. Phylogenetic characterization of two novel commensal bacteria involved with innate immune homeostasis in *Drosophila melanogaster*. Appl. Environ. Microbiol. 74, 6171–6177.

Sasser, M. 1990. Identification of bacteria by gas chromatography of cellular fatty acids. USFCC Newsletter. 20:16.

Shimwell, J. L., Carr, J. G., and Rhodes, M. E. 1960. Differentiation of *Acetomonas* and *Pseudomonas*. Microbiol. 23, 283–286.

Simão, F. A., Waterhouse, R. M., Ioannidis, P., Kriventseva, E. V., and Zdobnov, E. M. 2015. BUSCO: Assessing genome assembly and annotation completeness with single-copy orthologs. Bioinformatics. 31, 3210–3212.

Spitaels, F., Li, L., Wieme, A., Balzarini, T., Cleenwerck, I., Van Landschoot, A., De Vuyst, L., and Vandamme, P. 2014. *Acetobacter lambici* sp. nov., isolated from fermenting lambic beer. Int. J. Syst. Evol. Microbiol. 64, 1083–1089.

Tatusova, T., DiCuccio, M., Badretdin, A., Chetvernin, V., Nawrocki, E. P., Zaslavsky, L., Lomsadze, A., Pruitt, K. D., Borodovsky, M., and Ostell, J. 2016. NCBI prokaryotic genome annotation pipeline. Nucleic Acids Res. gkw569.

Tindall, B. J. 1990a. A comparative study of the lipid composition of *Halobacterium saccharovorum* from various sources. Syst. Appl. Microbiol. 13, 128–130.

Tindall, B. J. 1990b. Lipid composition of *Halobacterium lacusprofundi*. FEMS Microbiol. Lett. 66, 199–202.

Tittsler, R. P. and Sandholzer, L. A. 1936. The use of semi-solid agar for the detection of bacterial motility. J. Bacteriol. 31, 575.

Way, M. J. and Khoo, K. C. 1989. Relationships between *Helopeltis theobromae* damage and ants with special reference to Malaysian cocoa smallholdings. J. Plant Prot. Tropics. 6, 1–11.

Yamada, Y. 2016. Systematics of acetic acid bacteria, p. 1–50. Acetic acid bacteria, Springer, Tokyo, Japan.

Yamada, Y., Aida, K., and Uemura, T. 1968. Distribution of ubiquinone 10 and 9 in acetic acid bacteria and its relation to the classification of genera *Gluconobacter* and *Acetobacter*, especially of so-called intermediate strains. Agric. Biol. Chem. 32, 786–788.

Yamada, Y., Hosono, R., Lisdyanti, P., Widyastuti, Y., Saono, S., Uchimura, T., and Komagata, K. 1999. Identification of acetic acid bacteria isolated from Indonesian sources, especially of isolates classified in the genus *Gluconobacter*. J. Gen. Appl. Microbiol. 45, 23–28.

Yamada, Y. and Yukphan, P. 2008. Genera and species in acetic acid bacteria. Int. J. Food Microbiol. 125, 15–24.

Yarza, P., Yilmaz, P., Pruesse, E., Glöckner, F. O., Ludwig, W., Schleifer, K.-H., Whitman, W. B., Euzéby, J., Amann, R., and Rosselló-Móra, R. 2014. Uniting the classification of cultured and uncultured bacteria and archaea using 16S rRNA gene sequences. Nat. Rev. Microbiol. 12, 635–645.

Yong, H.-S., Song, S.-L., Chua, K.-O., Lim, P.-E., and Eamsobhana, P. 2019. Microbiota and potential opportunistic pathogens associated with male and female fruit flies of Malaysian *Bactrocera carambolae* (Insecta: Tephritidae). Meta Gene. 19, 185–192.

Yoon, S.-H., Ha, S.-M., Kwon, S., Lim, J., Kim, Y., Seo, H., and Chun, J. 2017. Introducing EzBioCloud: a taxonomically united database of 16S rRNA gene sequences and whole-genome assemblies. Int. J. Syst. Evol. Microbiol. 67, 1613–1617.

Yukphan, P., Malimas, T., Muramatsu, Y., Potacharoen, W., Tanasupawat, S., Nakagawa, Y., Tanticharoen, M., and Yamada, Y. 2011. *Neokomagataea* gen. nov., with descriptions of *Neokomagataea thailandica* sp. nov. and *Neokomagataea tanensis* sp. nov., osmotolerant acetic acid bacteria of the α-Proteobacteria. Biosci. Biotechnol. Biochem. 75, 419–426.

Yun, J.-H., Lee, J.-Y., Hyun, D.-W., Jung, M.-J., and Bae, J.-W. 2017. *Bombella apis* sp. nov., an acetic acid bacterium isolated from the midgut of a honey bee. Int. J. Syst. Evol. Microbiol. 67, 2184–2188.

