## Supplementary Table S1, S2, S3, S4, S5, S6; Supplementary Fig. S1, S2, S3, S4, S5, S6, S7, S8, S9, S10 for "*Oecophyllibacter saccharovorans* gen. nov. sp. nov., a bacterial symbiont of the weaver ant *Oecophylla smaragdina*"

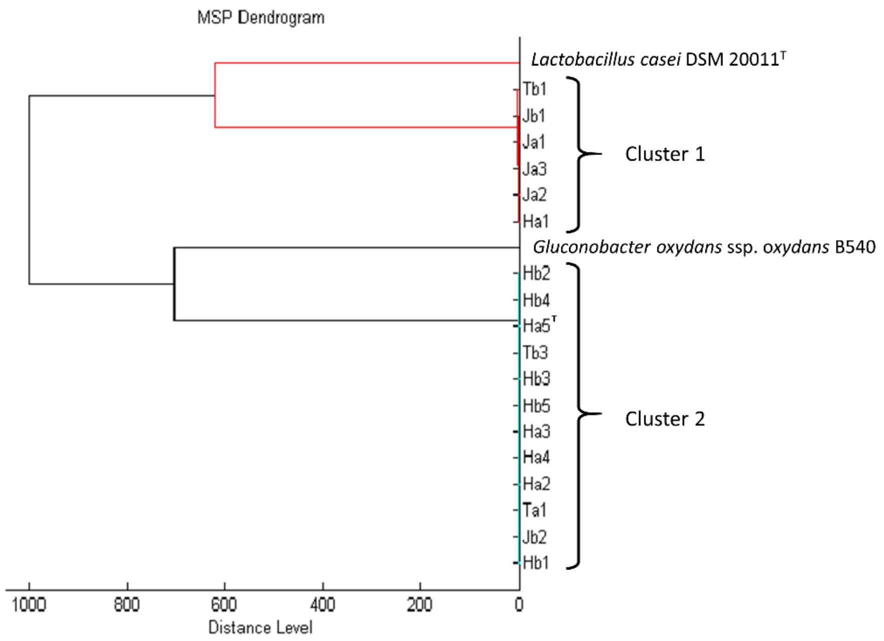


**Supplementary data Fig. S1. Main Spectrum Profile (MSP)-based dendrogram of isolates obtained from *O. smaragdina* in this study.**

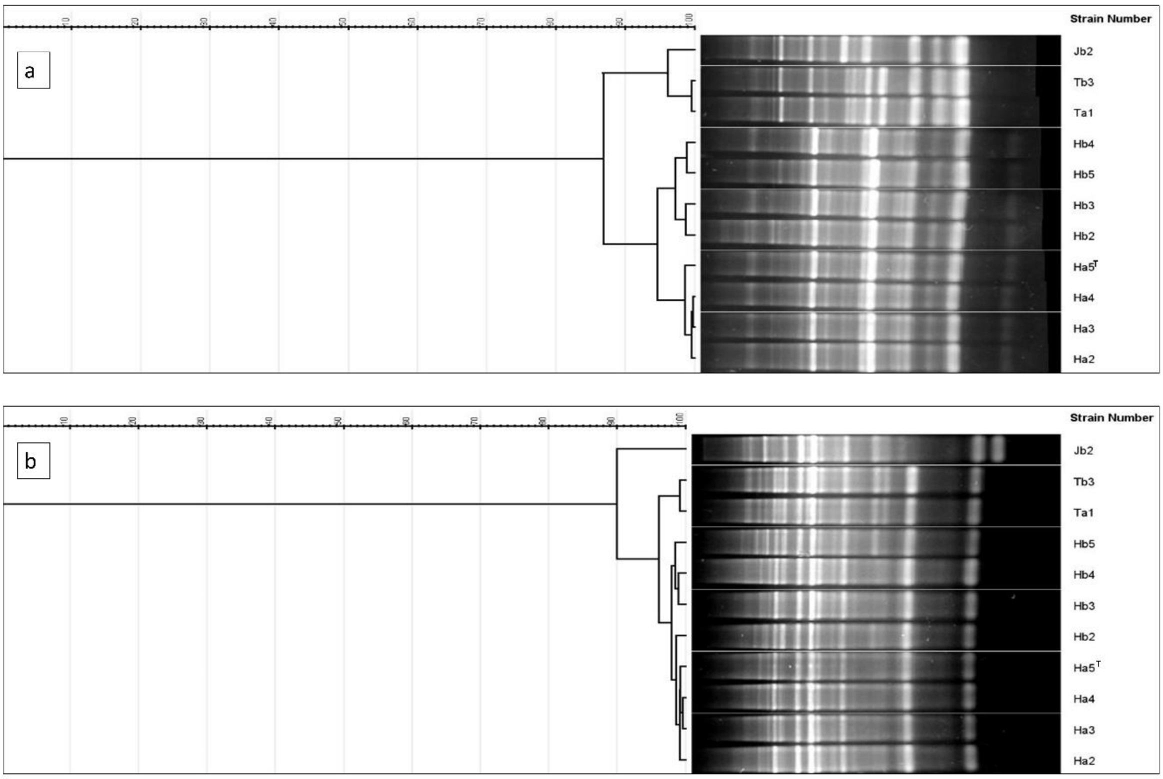


**Supplementary data Fig. S2. RAPD fingerprints differentiating strains in MALDI-TOF cluster 2 by (a) RAPD primer 270; (b) RAPD primer 272.** Cluster analysis was performed with GelJ by UPGMA method with similarity matrix calculated with the Pearson correlation.

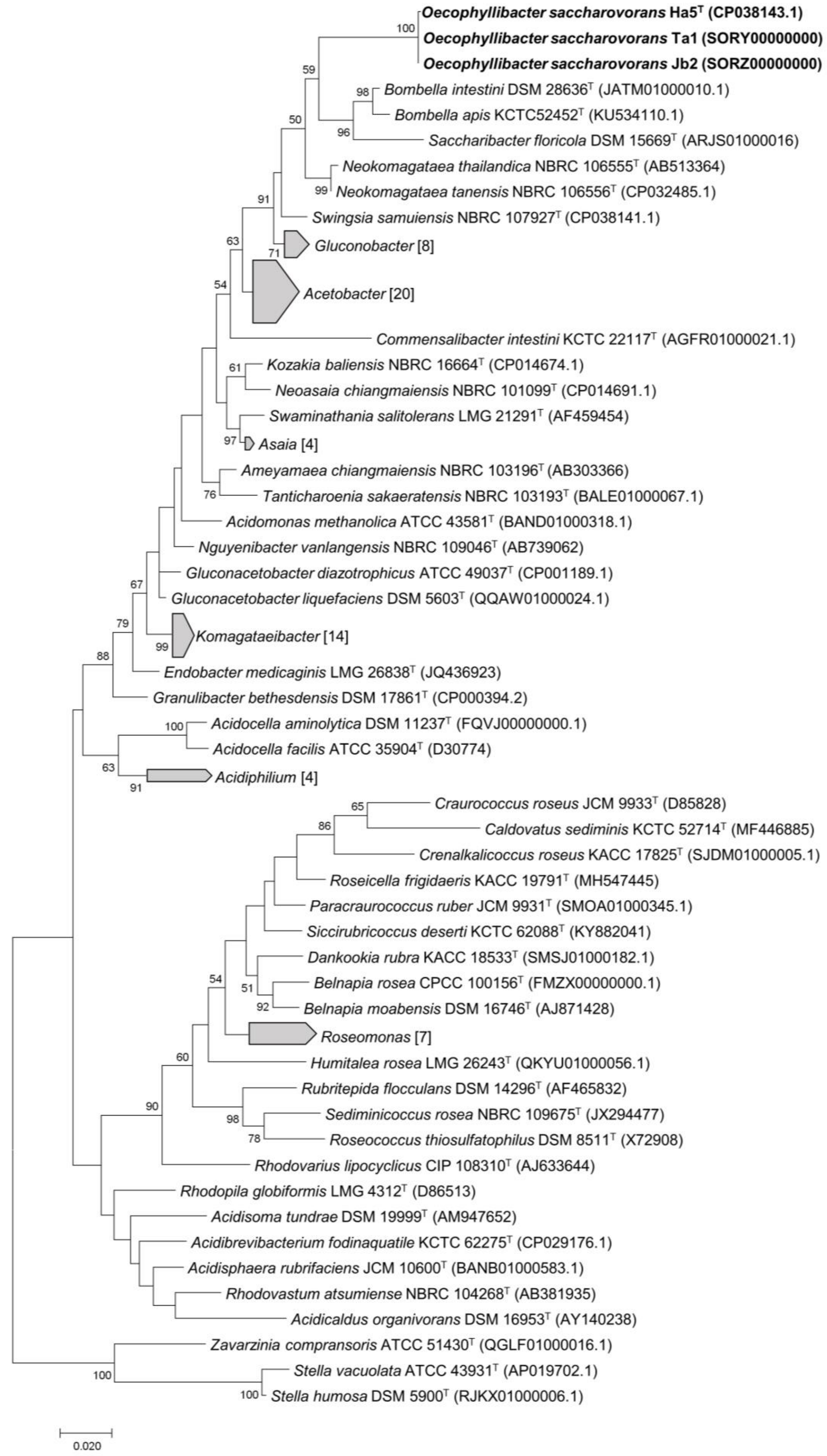


**Supplementary data Fig. S3. Maximum likelihood tree showing the phylogenetic relationship of strains Ha5$^{T}$, Ta1, Jb2 and their closest relatives in the family *Acetobacteraceae*.** The tree was constructed based on 16S rRNA gene sequences. Number displayed at nodes are bootstrap values, based on 1000 replications. Numbers in brackets represent the number of type strains included in the respective clusters. The tree was rooted with genera *Stella* and *Zavarzinia* as outgroups.

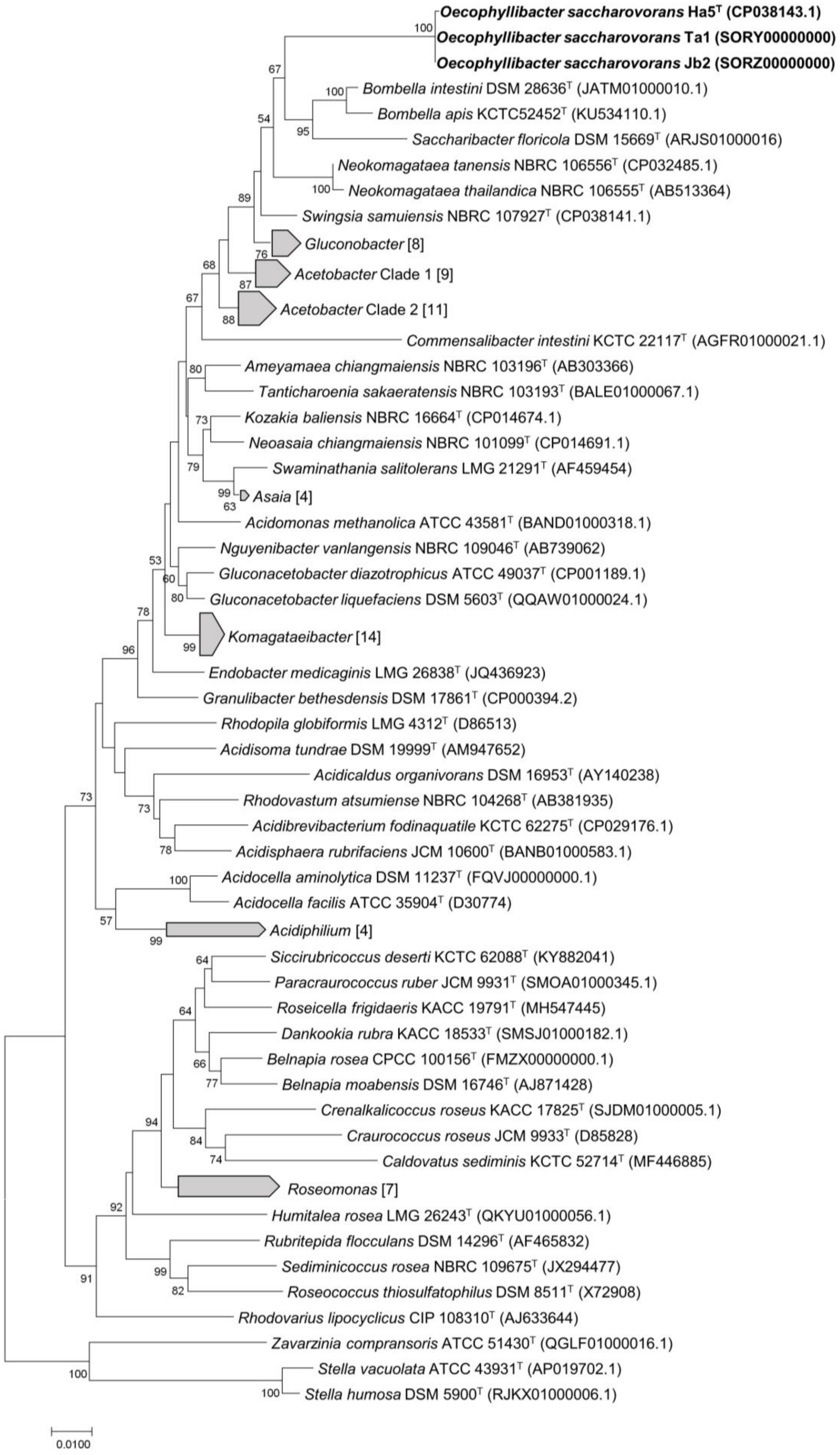


**Supplementary data Fig. S4. Neighbour-joining tree showing the relationship of strains Ha5$^T$, Ta1, Jb2 and their closest relatives in the family *Acetobacteraceae*.** The phylogenetic tree was constructed based on 16S rRNA gene sequences. Number displayed at nodes are bootstrap values, based on 1000 replications. Numbers in brackets represent the number of type strains included in the respective clusters. The tree was rooted using genera *Stella* and *Zavarzinia* as outgroup.

**Supplementary data Fig. S5. Average Nucleotide Identity (ANI)**

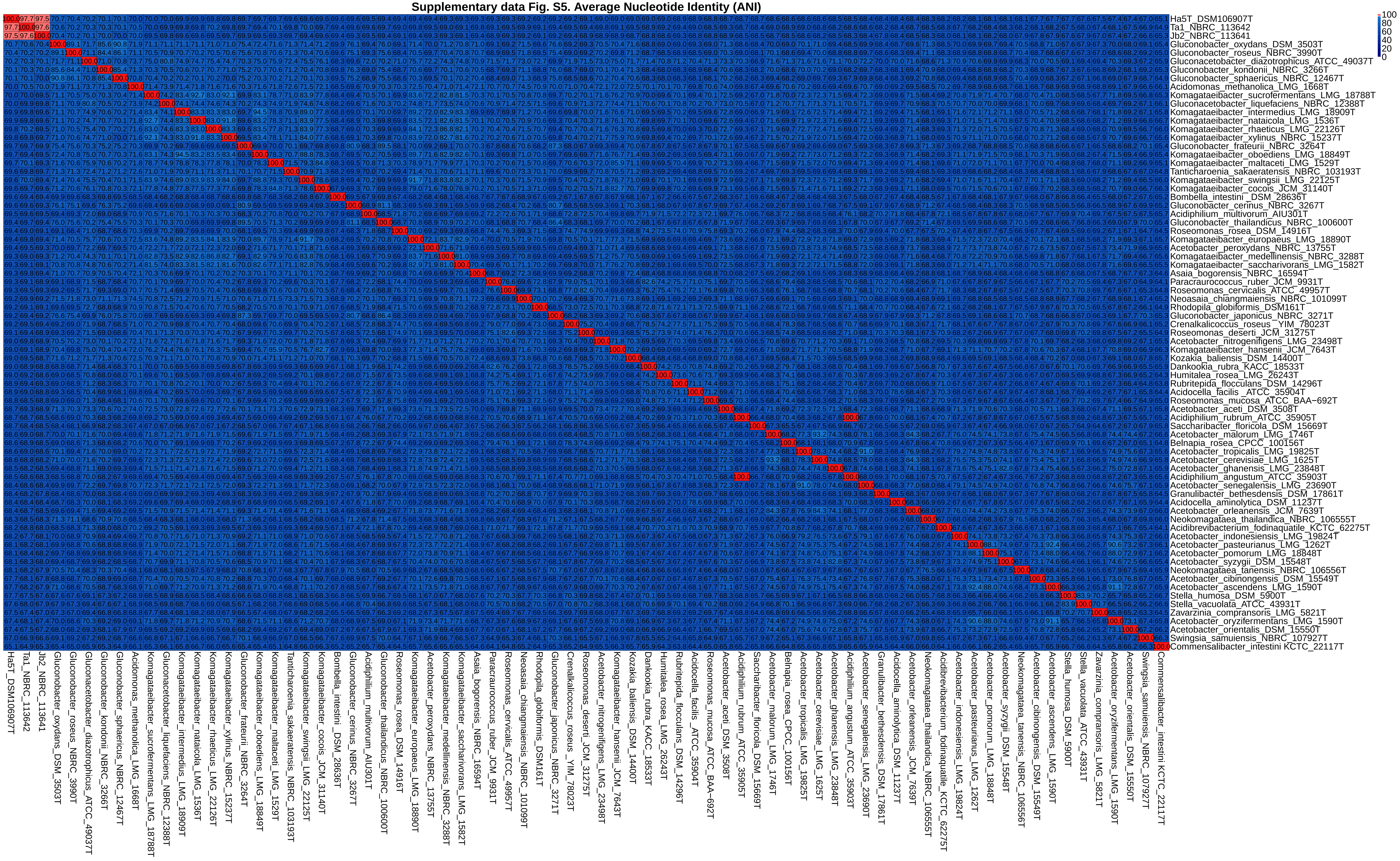

### Supplementary data Fig. S6. in silico DNA−DNA hybridization (DDH)

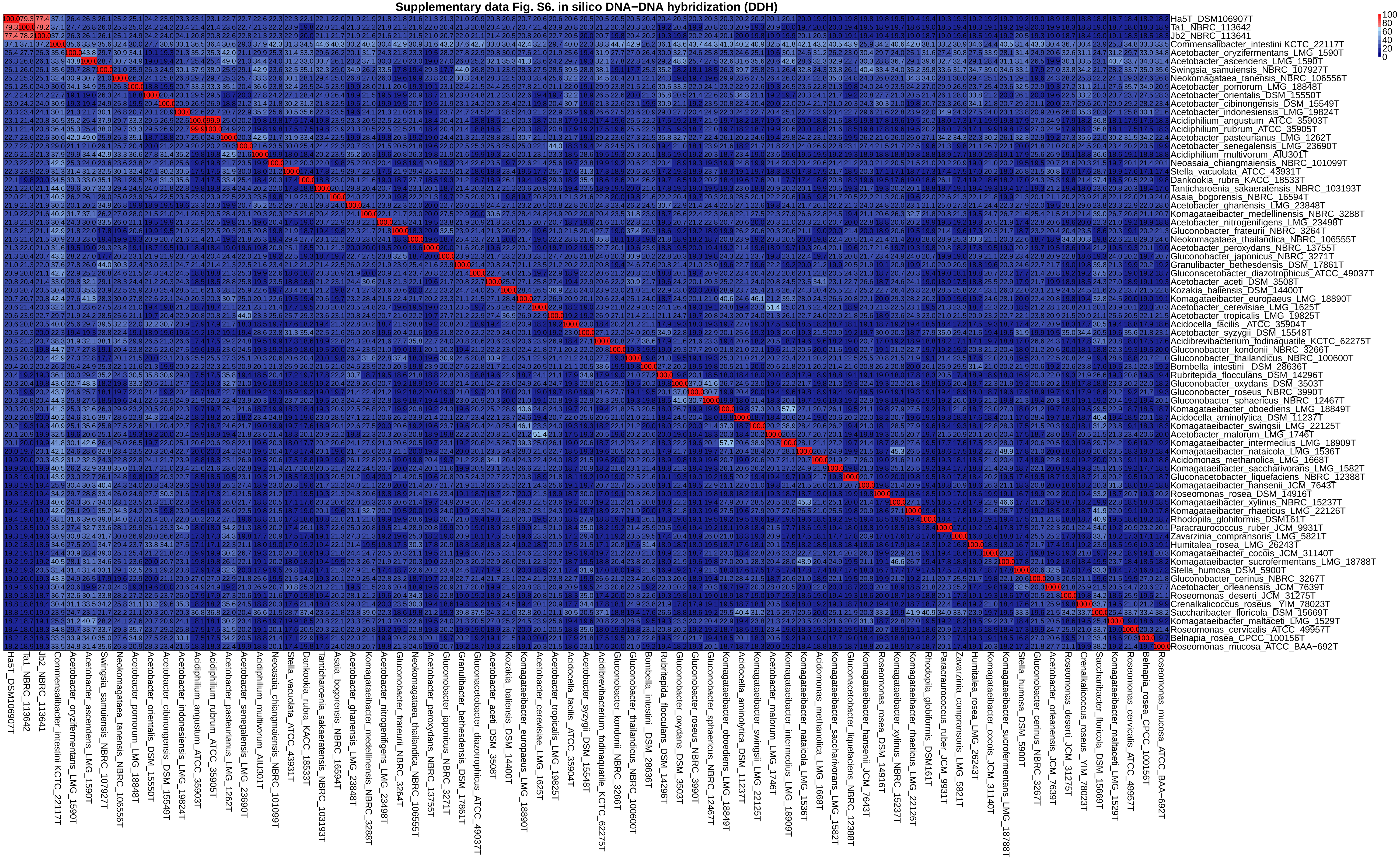

**Supplementary data Fig. S7. Average Amino acid Identity (AAI)**

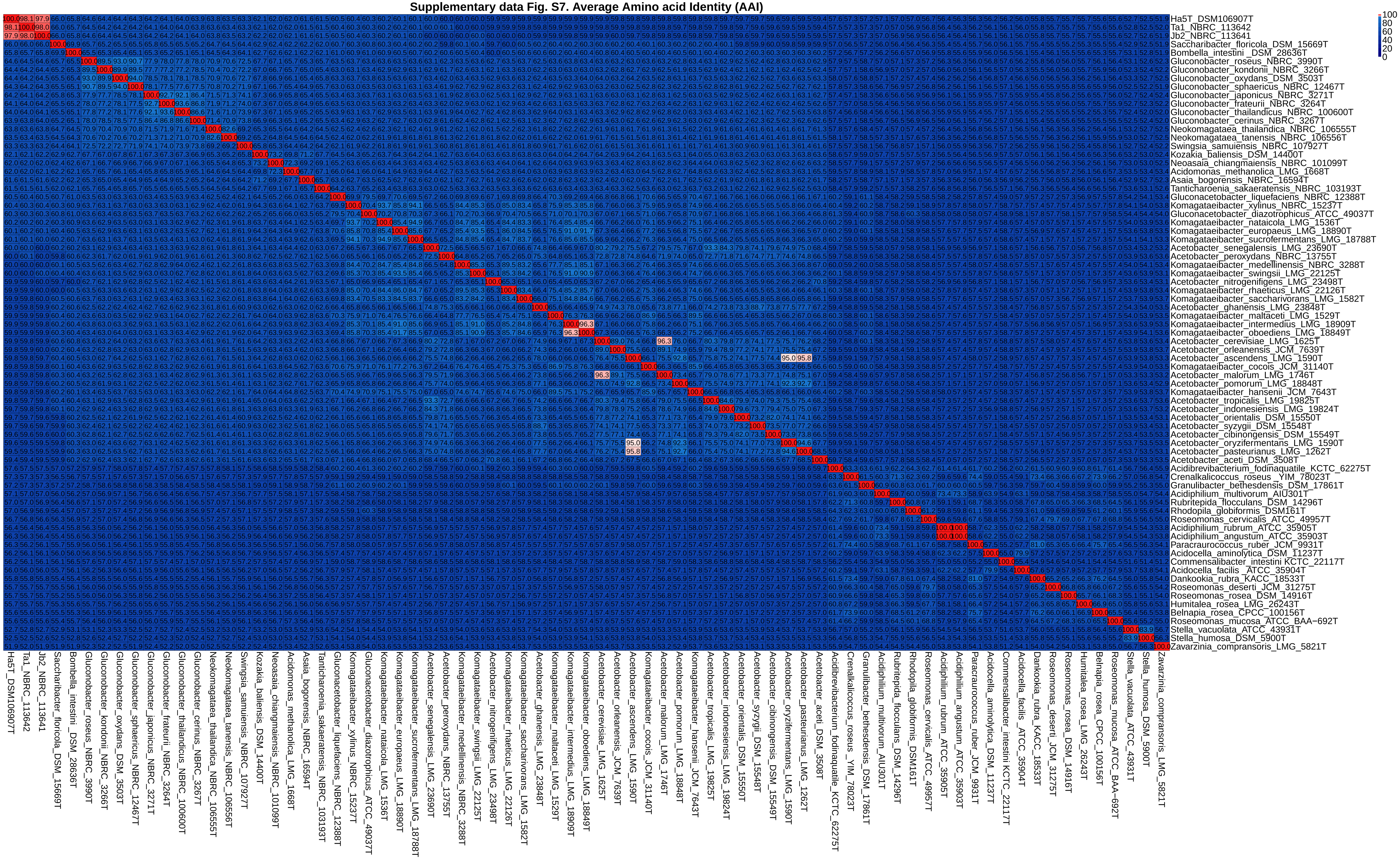

### Supplementary data Fig. S8. Percentage of Conserved Proteins (POCP)

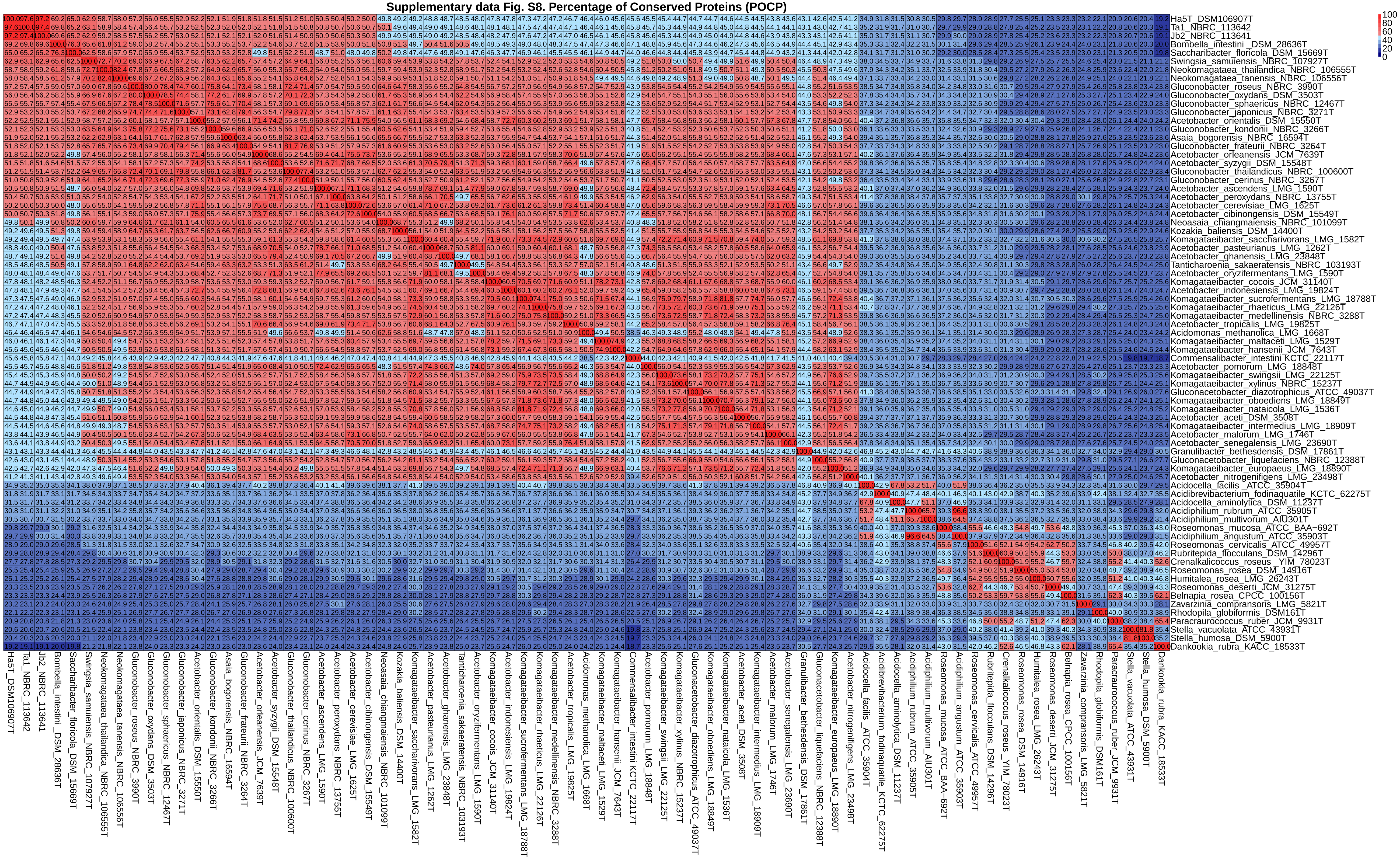

**Supplementary data Fig. S5. Average nucleotide identity (ANI) of strains Ha5$^T$, Ta1, Jb2 and their closest relatives in the family *Acetobacteraceae*.** ANI values lesser than 95% are colored in blue, indicating that the strains are of different species. Color intensity fades as the ANI values approach 95%. ANI values equal or greater than 95% are colored in red, indicating that these strains belong to the same species. Color intensity strengthens as the ANI values approach 100%.

**Supplementary data Fig. S6. *in silico* DNA–DNA hybridization (DDH) of strains Ha5$^T$, Ta1, Jb2 and their closest relatives in the family *Acetobacteraceae*.** DDH values lesser than 70% are colored in blue, indicating that the strains are of different species. Color intensity fades as the DDH values approach 70%. DDH values equal or greater than 70% are colored in red, indicating that these strains belong to the same species. Color intensity strengthens as the DDH values approach 100%.

**Supplementary data Fig. S7. Average amino acid identity (AAI) of strains Ha5$^T$, Ta1, Jb2 and their closest relatives in the family *Acetobacteraceae*.** AAI values lesser than 95% are colored in blue, indicating that the strains are of different genera. Color intensity fades as the AAI values approach 95%. AAI values equal or greater than 95% are colored in red, indicating that these strains belong to the same genus. Color intensity strengthens as the AAI values approach 100%.

**Supplementary data Fig. S8. Percentage of Conserved Proteins (POCP) of strains Ha5$^T$, Ta1, Jb2 and their closest relatives in the family *Acetobacteraceae*.** POCP values lesser than 50% are colored in blue, indicating that the strains are of different species. Color intensity fades as the POCP values approach 50%. POCP values equal or greater than 50% are colored in red, indicating that these strains belong to the same species. Color intensity strengthens as the POCP values approach 100%.

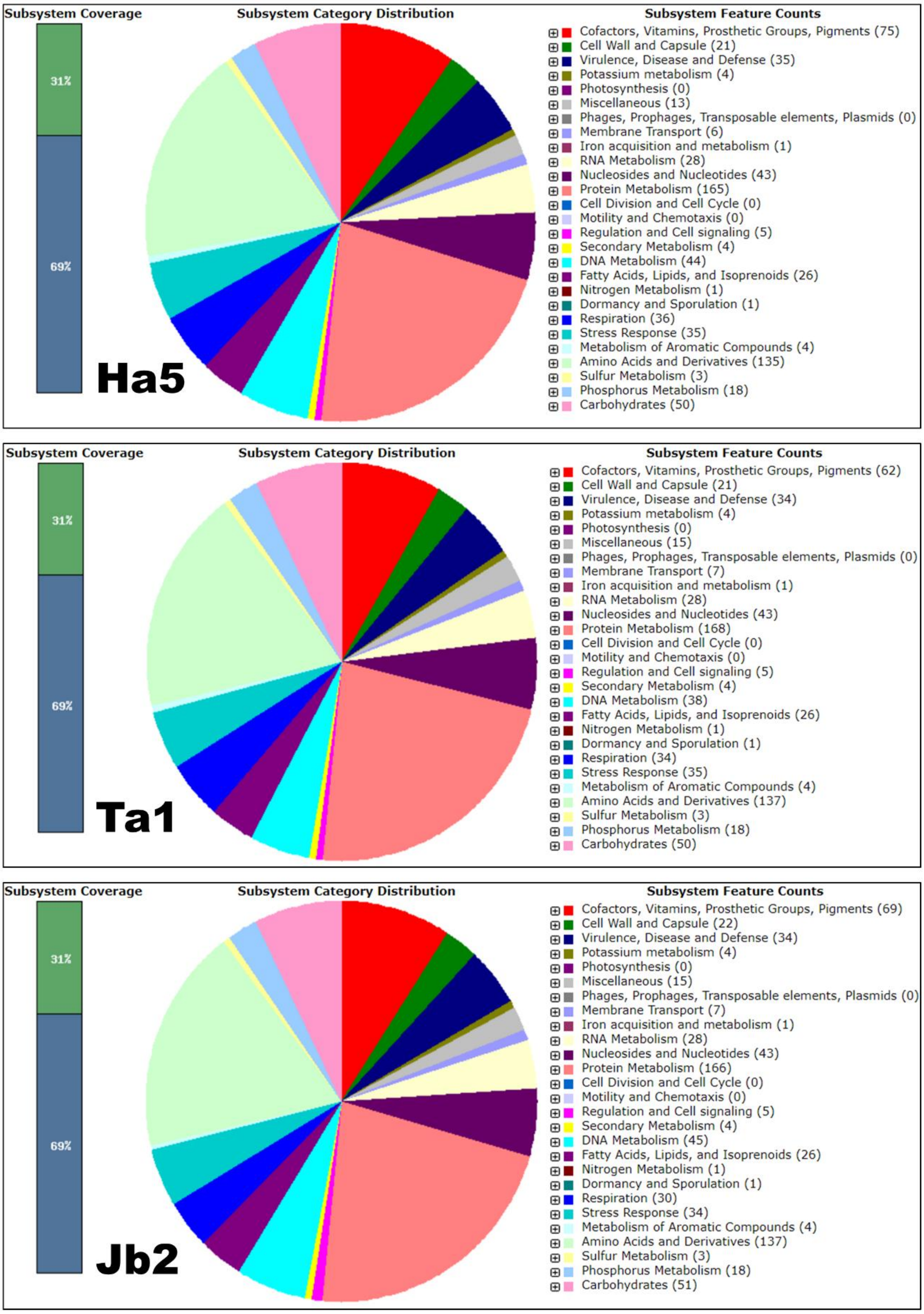


**Supplementary data Fig. S9.** Gene distribution of strains Ha5$^{T}$, Ta1 and Jb2 into RAST subsystem categories.

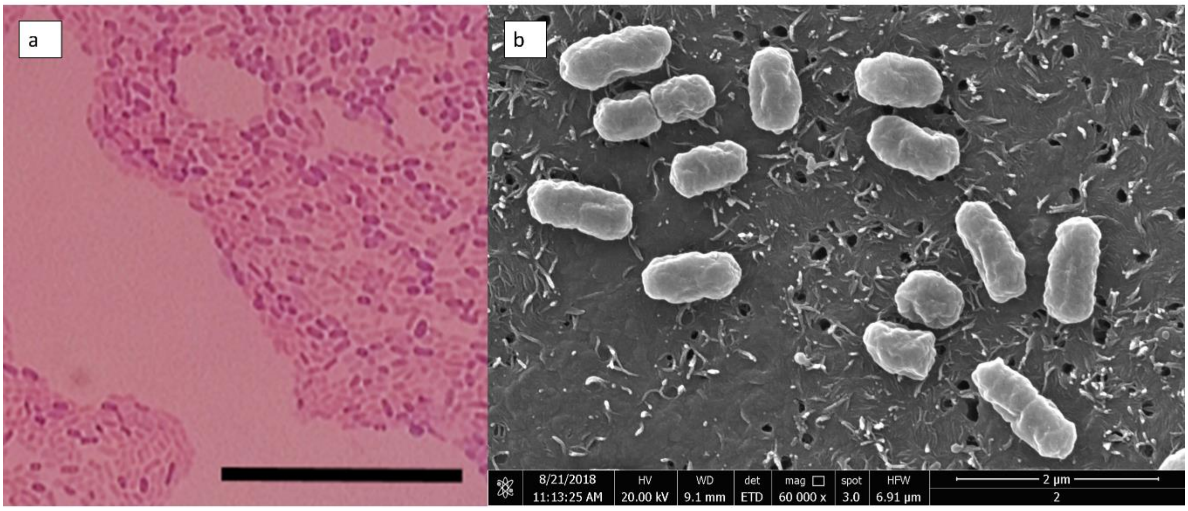


**Supplementary data Fig. S10.** Cellular morphology of strain Ha5$^T$. (a) Gram-stained cells of Ha5$^T$ under light microscope. Bar, 10 µm. (b) Scanning electron micrograph (SEM) showing the rod-shaped cells of strain Ha5$^T$. Bar, 2 µm.

**Supplementary data Table S1.** List of *O. smaragdina* colonies sampled for cultivation of bacteria from family *Acetobacteraceae*.

| Colony | Location | Coordinates | Collection date |
|---|---|---|---|
| H | Kuala Lumpur, University of Malaya | 3°07'21.9"N 101°39'22.7"E | 14 June 2017 |
| T | Selangor, Petaling Jaya | 3°08'50.8"N 101°37'55.3"E | 11 July 2017 |
| J | Selangor, Petaling Jaya | 3°06'14.4"N 101°38'56.6"E | 11 July 2017 |

**Supplementary data Table S2.** List of genomes of *Acetobacteraceae* type members included in this study.

| Strains | Genome accession | No. of contig | Status | Genome size (Mb) | G+C content | Number of gene | Protein coding genes | BUSCO completeness (%) |
|---|---|---|---|---|---|---|---|---|
| *Acetobacter aceti* DSM 3508$^{T}$ | SLZP00000000.1 | 57 | Draft | 3.63 | 57.1 | 3376 | 3200 | 97.7 |
| *Acidocella facilis* ATCC 35904$^{T}$ | JHYG00000000.1 | 48 | Draft | 3.40 | 64.5 | 3272 | 3139 | 97.7 |
| *Acidomonas methanolica* LMG 1668$^{T}$ | BAND00000000.1 | 546 | Draft | 3.69 | 64.6 | 3510 | 3307 | 96.0 |
| *Asaia bogorensis* NBRC 16594$^{T}$ | AP014690.1 | 1 | Complete | 3.20 | 59.8 | 2803 | 2692 | 97.3 |
| *Bombella intestini* DSM 28636$^{T}$ | JATM00000000.1 | 12 | Draft | 2.02 | 55.0 | 1632 | 1574 | 96.4 |
| *Commensalibacter intestini* KCTC 22117$^{T}$ | AGFR00000000.1 | 26 | Draft | 2.45 | 36.8 | 2255 | 2160 | 97.3 |
| *Crenalkalicoccus roseus* YIM 78023$^{T}$ | SJDM00000000.1 | 127 | Draft | 4.43 | 73.9 | 4231 | 4037 | 96.4 |
| *Dankookia rubra* KACC 18533$^{T}$ | SMSJ00000000.1 | 458 | Draft | 7.78 | 70.1 | 7451 | 7102 | 95.0 |
| *Gluconacetobacter liquefaciens* NBRC 12388$^{T}$ | QQAW00000000.1 | 31 | Draft | 4.18 | 64.4 | 3700 | 3511 | 97.3 |
| *Gluconobacter oxydans* DSM 3503$^{T}$ | SMCY00000000.1 | 88 | Draft | 2.92 | 60.8 | 2782 | 2605 | 97.8 |
| *Granulibacter bethesdensis* DSM 17861$^{T}$ | CP000394.2 | 1 | Complete | 2.71 | 59.1 | 2471 | 2384 | 97.7 |
| Ha5 = DSM106907$^{T}$ | CP038143.1 | 2 | Complete | 1.95 | 61.6 | 1689 | 1617 | 97.3 |
| *Humitalea rosea* LMG 26243$^{T}$ | QKYU00000000.1 | 73 | Draft | 4.97 | 69.6 | 4723 | 4668 | 97.3 |
| Jb2 = NBRC 113641 | SORZ00000000 | 3 | Draft | 1.94 | 61.6 | 1680 | 1612 | 97.7 |
| *Komagataeibacter xylinus* NBRC 15237$^{T}$ | BCTP00000000.1 | 156 | Draft | 3.63 | 62.3 | 3441 | 3197 | 97.3 |
| *Kozakia baliensis* DSM 14400$^{T}$ | CP014674.1 | 7 | Complete | 3.51 | 57.4 | 3381 | 3130 | 96.8 |
| *Neoasaia chiangmaiensis* NBRC 101099$^{T}$ | CP014691.1 | 1 | Complete | 3.41 | 61.5 | 3210 | 3051 | 97.7 |
| *Neokomagataea tanensis* NBRC 106556$^{T}$ | CP032485.1 | 2 | Complete | 2.58 | 52.0 | 2351 | 2155 | 95.9 |
| *Neokomagataea thailandica* NBRC 106555$^{T}$ | BCZB00000000.1 | 83 | Draft | 2.49 | 52.4 | 2317 | 2156 | 96.8 |
| *Paracraurococcus ruber* JCM 9931$^{T}$ | SMOA00000000.1 | 786 | Draft | 7.17 | 72.8 | 6537 | 6352 | 95.5 |
| *Rhodopila globiformis* DSM161$^{T}$ | NHRY00000000.1 | 278 | Draft | 7.25 | 67.1 | 6419 | 6196 | 97.3 |
| *Rubritepida flocculans* DSM 14296$^{T}$ | AUDH00000000.1 | 74 | Draft | 3.83 | 73.4 | 3742 | 3637 | 97.8 |
| *Saccharibacter floricola* DSM 15669$^{T}$ | ARJS00000000.1 | 43 | Draft | 2.38 | 51.2 | 2264 | 2133 | 97.3 |
| *Stella humosa* DSM 5900$^{T}$ | RJKX00000000.1 | 20 | Draft | 5.83 | 69.9 | 5467 | 5365 | 98.2 |
| *Swingsia samuiensis* NBRC 107927$^{T}$ | CP038141.1 | 2 | Complete | 2.19 | 45.1 | 2098 | 1988 | 98.6 |
| Ta1 = NBRC 113642 | SORY00000000 | 4 | Draft | 1.92 | 61.7 | 1673 | 1601 | 97.3 |
| *Tanticharoenia sakaeratensis* NBRC 103193$^{T}$ | BALE00000000.1 | 94 | Draft | 3.51 | 64.2 | 3137 | 3051 | 97.8 |
| *Zavarzinia compransoris* LMG 5821$^{T}$ | QGLF00000000.1 | 17 | Draft | 4.76 | 68.1 | 4521 | 4401 | 98.7 |
| *Acidibrevibacterium fodinaquatile* KCTC 62275$^{T}$ | CP029176.1 | 4 | Complete | 4.04 | 65.5 | 3826 | 3633 | 97.7 |
| *Acetobacter ascendens* LMG 1590$^{T}$ | CP015164.1 | 4 | Complete | 3.00 | 53.2 | 2938 | 2560 | 97.7 |
| *Acetobacter cerevisiae* LMG 1625$^{T}$ | LHZA00000000.1 | 157 | Draft | 3.09 | 58.0 | 2832 | 2658 | 96.8 |
| *Acetobacter cibinongensis* DSM 15549$^{T}$ | BAMV00000000.1 | 120 | Draft | 3.11 | 54.4 | 2759 | 2646 | 96.8 |
| *Acetobacter ghanensis* LMG 23848$^{T}$ | LN609302.1 | 3 | Complete | 2.84 | 56.9 | 2686 | 2454 | 94.6 |
| *Acetobacter indonesiensis* LMG 19824$^{T}$ | BAMW00000000.1 | 181 | Draft | 3.42 | 53.9 | 3068 | 2917 | 98.2 |
| *Acetobacter malorum* LMG 1746$^{T}$ | LHZC00000000.1 | 57 | Draft | 3.83 | 56.7 | 3454 | 3299 | 97.3 |
| *Acetobacter nitrogenifigens* LMG 23498$^{T}$ | AUBI00000000.1 | 51 | Draft | 4.27 | 60.5 | 3802 | 3613 | 97.3 |
| *Acetobacter orientalis* DSM 15550$^{T}$ | BAMX00000000.1 | 66 | Draft | 2.91 | 52.4 | 2560 | 2469 | 97.7 |

| | | | | | | | | |
|---|---|---|---|---|---|---|---|---|
| *Acetobacter orleanensis* JCM 7639$^T$ | BAMY00000000.1 | 269 | Draft | 3.01 | 56.4 | 2715 | 2564 | 97.3 |
| *Acetobacter oryzifermentans* LMG 1590$^T$ | CP011120.1 | 4 | Complete | 3.10 | 52.4 | 3009 | 2787 | 97.3 |
| *Acetobacter pasteurianus* LMG 1262$^T$ | CADO00000000.1 | 141 | Draft | 2.98 | 53.1 | 2883 | 2698 | 97.3 |
| *Acetobacter peroxydans* NBRC 13755$^T$ | BJMV00000000.1 | 29 | Draft | 2.70 | 60.5 | 2475 | 2379 | 96.8 |
| *Acetobacter pomorum* LMG 18848$^T$ | PEBQ00000000.1 | 212 | Draft | 3.33 | 51.7 | 3168 | 2973 | 97.3 |
| *Acetobacter senegalensis* LMG 23690$^T$ | LHZU00000000.1 | 148 | Draft | 3.93 | 55.6 | 3492 | 3293 | 97.8 |
| *Acetobacter syzygii* DSM 15548$^T$ | BAMZ00000000.1 | 94 | Draft | 2.69 | 55.5 | 2478 | 2365 | 95.5 |
| *Acetobacter tropicalis* LMG 19825$^T$ | LHZQ00000000.1 | 222 | Draft | 3.56 | 55.8 | 3194 | 2962 | 97.7 |
| *Acidiphilium angustum* ATCC 35903$^T$ | JNJH00000000.1 | 206 | Draft | 4.18 | 63.6 | 3921 | 3730 | 97.3 |
| *Acidiphilium multivorum* AIU301$^T$ | AP012035.1 | 9 | Complete | 4.21 | 67.0 | 3991 | 3803 | 98.2 |
| *Acidiphilium rubrum* ATCC 35905$^T$ | FTNE00000000.1 | 78 | Draft | 3.98 | 63.7 | 3752 | 3692 | 97.3 |
| *Acidocella aminolytica* DSM 11237$^T$ | FQVJ00000000.1 | 109 | Draft | 3.96 | 58.9 | 3931 | 3685 | 97.8 |
| *Belnapia rosea* CPCC 100156$^T$ | FMZX00000000.1 | 119 | Draft | 5.99 | 69.8 | 5723 | 5467 | 97.7 |
| *Gluconacetobacter diazotrophicus* ATCC 49037$^T$ | CP001189.1 | 2 | Complete | 3.91 | 66.3 | 3647 | 3457 | 99.1 |
| *Gluconobacter cerinus* NBRC 3267$^T$ | BEWM00000000.1 | 49 | Draft | 3.59 | 55.6 | 3310 | 3118 | 97.7 |
| *Gluconobacter frateurii* NBRC 3264$^T$ | BEWN00000000.1 | 10 | Draft | 3.31 | 56.1 | 3095 | 2976 | 97.3 |
| *Gluconobacter japonicus* NBRC 3271$^T$ | BEWO00000000.1 | 75 | Draft | 3.16 | 56.1 | 2995 | 2839 | 97.7 |
| *Gluconobacter kondonii* NBRC 3266$^T$ | BEWP00000000.1 | 63 | Draft | 3.27 | 58.3 | 3140 | 2950 | 96.8 |
| *Gluconobacter roseus* NBRC 3990$^T$ | BJLY00000000.1 | 12 | Draft | 2.88 | 59.9 | 2643 | 2504 | 97.3 |
| *Gluconobacter sphaericus* NBRC 12467$^T$ | BJMK00000000.1 | 97 | Draft | 3.06 | 58.2 | 2887 | 2837 | 97.3 |
| *Gluconobacter thailandicus* NBRC 100600$^T$ | BANH00000000.1 | 191 | Draft | 3.44 | 56.2 | 3209 | 3038 | 97.3 |
| *Komagataeibacter cocois* JCM 31140$^T$ | QEXL00000000.1 | 75 | Draft | 3.41 | 62.3 | 3117 | 2934 | 95.9 |
| *Komagataeibacter europaeus* LMG 18890$^T$ | CADP00000000.1 | 321 | Draft | 4.23 | 61.3 | 4229 | 3622 | 96.4 |
| *Komagataeibacter hansenii* JCM 7643$^T$ | BANE00000000.1 | 467 | Draft | 3.71 | 59.3 | 3324 | 3149 | 96.4 |
| *Komagataeibacter intermedius* LMG 18909$^T$ | BANF00000000.1 | 943 | Draft | 3.88 | 61.6 | 3656 | 3430 | 97.7 |
| *Komagataeibacter maltaceti* LMG 1529$^T$ | NKUE00000000.1 | 127 | Draft | 3.64 | 63.2 | 3347 | 3116 | 97.3 |
| *Komagataeibacter medellinensis* NBRC 3288$^T$ | AP012159.1 | 8 | Complete | 3.51 | 60.6 | 3378 | 3078 | 96.8 |
| *Komagataeibacter nataicola* LMG 1536$^T$ | NIRT00000000.1 | 106 | Draft | 3.67 | 61.5 | 3526 | 3274 | 97.7 |
| *Komagataeibacter oboediens* LMG 18849$^T$ | NKTX00000000.1 | 278 | Draft | 3.78 | 61.4 | 3722 | 3395 | 97.7 |
| *Komagataeibacter rhaeticus* LMG 22126$^T$ | NKTZ00000000.1 | 64 | Draft | 3.47 | 63.5 | 3175 | 2984 | 97.3 |
| *Komagataeibacter saccharivorans* LMG 1582$^T$ | NKTY00000000.1 | 107 | Draft | 3.35 | 61.6 | 3103 | 2889 | 97.3 |
| *Komagataeibacter sucrofermentans* LMG 18788$^T$ | NKUA00000000.1 | 83 | Draft | 3.36 | 62.3 | 3150 | 2979 | 97.3 |
| *Komagataeibacter swingsii* LMG 22125$^T$ | NKUB00000000.1 | 113 | Draft | 3.73 | 62.4 | 3497 | 3292 | 97.7 |
| *Roseomonas cervicalis* ATCC 49957$^T$ | ADVL00000000.1 | 498 | Draft | 5.10 | 72.0 | 4615 | 3821 | 93.7 |
| *Roseomonas deserti* JCM 31275$^T$ | MLCO00000000.1 | 597 | Draft | 6.25 | 71.1 | 5978 | 5405 | 93.7 |
| *Roseomonas mucosa* ATCC BAA-692$^T$ | JHWD00000000.1 | 77 | Draft | 4.86 | 70.4 | 4461 | 4259 | 97.3 |
| *Roseomonas rosea* DSM 14916$^T$ | FQZF00000000.1 | 81 | Draft | 5.34 | 70.7 | 5092 | 5027 | 96.4 |
| *Stella vacuolata* ATCC 43931$^T$ | AP019702.1 | 1 | Complete | 5.83 | 70.9 | 5517 | 5371 | 98.2 |
| *Acetobacter okinawensis* JCM 25146$^T$ | BAJU00000000.1 | 127 | Draft | 3.17 | 57.6 | 3062 | 2271 | 76.9[1] |
| *Acetobacter papayae* JCM 25143$^T$ | BAIN00000000.1 | 177 | Draft | 3.04 | 59.3 | 2964 | 2032 | 69.2[1] |
| *Acetobacter persici* JCM 25330$^T$ | BAJW00000000.1 | 236 | Draft | 3.70 | 57.2 | 3518 | 2521 | 75.2[1] |

| | | | | | | | | |
|---|---|---|---|---|---|---|---|---|
| *Acidisphaera rubrifaciens* JCM 10600$^{T}$ | BANB00000000.1 | 1239 | Draft | 3.86 | 70.3 | 3512 | 2788 | 71.9[1] |
| *Asaia astilbis* JCM 15831$^{T}$ | BAJT00000000.1 | 49 | Draft | 3.15 | 58.0 | 2969 | 2305 | 78.3[1] |
| *Asaia platycodi* JCM 25414$^{T}$ | BAKW00000000.1 | 35 | Draft | 3.15 | 59.3 | 2969 | 2085 | 71.1[1] |
| *Asaia prunellae* JCM 25354$^{T}$ | BAJV00000000.1 | 38 | Draft | 3.18 | 55.8 | 2941 | 2318 | 78.8[1] |
| *Komagataeibacter kakiaceti* JCM 25156$^{T}$ | BAIO00000000.1 | 947 | Draft | 3.13 | 62.1 | 3667 | 2121 | 49.4[1] |
| *Belnapia moabensis* DSM 16746$^{T}$ | JQKB00000000.1 | 282 | Draft | 6.73 | 68.9 | 6334[2] | 5826 | 96.4 |
| *Gluconacetobacter entanii* LMG 21788$^{T}$ | NKUF00000000.1 | 157 | Draft | 3.60 | 62.6 | 3375[2] | 3113 | 97.3 |
| *Roseomonas aerilata* DSM 19363$^{T}$ | JONP00000000.1 | 151 | Draft | 6.43 | 69.7 | 6007[2] | 5624 | 96.9 |
| Roseomonas *gilardii* subsp. *rosea* ATCC BAA-691$^{T}$ | JADY00000000.1 | 51 | Draft | 4.61 | 70.7 | 4238[2] | 4068 | 96.8 |
| *Roseomonas rhizosphaerae* KACC 17225$^{T}$ | PDNU00000000.1 | 114 | Draft | 4.65 | 71.9 | 4375[3] | 4205 | 97.3 |
| *Roseomonas stagni* DSM 19981$^{T}$ | FOSQ00000000.1 | 44 | Draft | 6.38 | 70.6 | 5,984[2] | 5,924 | 95.9 |

[1] Genome excluded from analysis due to low BUSCO genome completeness

[2] Genome excluded from analysis due to absence of 16S rRNA gene

[3] Genome excluded from analysis due to ambiguous 16S rRNA gene identity

**Supplementary data Table S3.** List of bacterial isolates cultured from different enrichment media

| Bacterial isolates | Sample ID | Enrichment medium | MALDI-TOF cluster |
|---|---|---|---|
| Ha1 | H | A | 1 |
| Ha2 | H | A | 2 |
| Ha3 | H | A | 2 |
| Ha4 | H | A | 2 |
| Ha5$^{T}$ | H | A | 2 |
| Hb1 | H | B | 2 |
| Hb2 | H | B | 2 |
| Hb3 | H | B | 2 |
| Hb4 | H | B | 2 |
| Hb5 | H | B | 2 |
| Ja1 | J | A | 1 |
| Ja2 | J | A | 1 |
| Ja3 | J | A | 1 |
| Jb1 | J | B | 1 |
| Jb2 | J | B | 2 |
| Ta1 | T | A | 2 |
| Tb1 | T | B | 1 |
| Tb3 | T | B | 2 |

**Supplementary data Table S4.** Results of preliminary 16S rRNA gene identification of selected isolates from each MALDI-TOF cluster with reference to EzBioCloud 16S database.

| Bacterial isolates | *O. smaragdina* colony | MALDI-TOF cluster | Top 16S rRNA gene match in EzBioCloud 16S database | Percentage identity (%) |
|---|---|---|---|---|
| Tb1 | T | 1 | *L. sanfranciscensis* ATCC 27651$^{T}$ | 98.15 |
| Jb1 | J | 1 | *L. sanfranciscensis* ATCC 27651$^{T}$ | 98.52 |
| Ha1 | H | 1 | *L. sanfranciscensis* ATCC 27651$^{T}$ | 98.52 |
| Ha5 | H | 2 | *N. tanensis* NBRC106556$^{T}$ | 94.23 |
| Ta1 | T | 2 | *N. tanensis* NBRC106556$^{T}$ | 94.31 |
| Jb2 | J | 2 | *N. tanensis* NBRC106556$^{T}$ | 94.31 |

**Supplementary data Table S5.** Pairwise 16S rRNA gene sequence similarity between novel strains Ha5$^{T}$, Ta1, Jb2 and representatives of closely related genera and species.

| Strains | Ha5$^{T}$ | Ta1 | Jb2 | *N. tanensis* NBRC 106556T | *B. intestine* DSM 28636T | *N. thailandica* NBRC 106555$^{T}$ | *S. samuiensis* NBRC 107927$^{T}$ | *G.. oxydans* DSM 3503$^{T}$ | *S. floricola* DSM 15669$^{T}$ | *A. chiangmaiensis* NBRC 103196$^{T}$ | *A. bogorensis* NBRC 16594$^{T}$ | *T. sakaeratensis* NBRC 103193$^{T}$ | *A. aceti* DSM 3508$^{T}$ |
|---|---|---|---|---|---|---|---|---|---|---|---|---|---|
| Ha5 DSM 106907$^{T}$ | 100 | 99.93 | 99.93 | 94.56 | 94.09 | 94.03 | 93.96 | 93.64 | 93.22 | 92.78 | 92.7 | 92.58 | 92.57 |
| Jb2 NBRC 113641 | 99.93 | 100 | 100 | 94.63 | 94.16 | 94.1 | 94.03 | 93.7 | 93.29 | 92.85 | 92.77 | 92.64 | 92.63 |
| Ta1 NBRC 113642 | 99.93 | 100 | 100 | 94.63 | 94.16 | 94.1 | 94.03 | 93.7 | 93.29 | 92.85 | 92.77 | 92.64 | 92.63 |
| *Neokomagataea tanensis* NBRC 106556$^{T}$ | 94.56 | 94.63 | 94.63 | 100 | 95.97 | 99.72 | 97.45 | 96.45 | 94.69 | 95.54 | 95.72 | 95.86 | 95.18 |
| *Bombella intestini* DSM 28636$^{T}$ | 94.09 | 94.16 | 94.16 | 95.97 | 100 | 95.45 | 95.64 | 95.51 | 96.24 | 94.06 | 94.78 | 93.99 | 95.11 |
| *Neokomagataea thailandica* NBRC 106555$^{T}$ | 94.03 | 94.1 | 94.1 | 99.72 | 95.45 | 100 | 97.02 | 95.95 | 94.24 | 95.26 | 95.18 | 95.34 | 94.61 |
| *Swingsia samuiensis* NBRC 107927$^{T}$ | 93.96 | 94.03 | 94.03 | 97.45 | 95.64 | 97.02 | 100 | 96.58 | 94.5 | 95.76 | 95.86 | 95.86 | 95.92 |
| *Gluconobacter oxydans* DSM 3503$^{T}$ | 93.64 | 93.7 | 93.7 | 96.45 | 95.51 | 95.95 | 96.58 | 100 | 94.37 | 95.54 | 95.85 | 96.12 | 95.85 |
| *Saccharibacter floricola* DSM 15669$^{T}$ | 93.22 | 93.29 | 93.29 | 94.69 | 96.24 | 94.24 | 94.5 | 94.36 | 100 | 93.35 | 93.59 | 93.24 | 93.97 |
| *Ameyamaea chiangmaiensis* NBRC 103196$^{T}$ | 92.78 | 92.85 | 92.85 | 95.54 | 94.06 | 95.26 | 95.76 | 95.54 | 93.35 | 100 | 97.03 | 98.02 | 96.24 |
| *Asaia bogorensis* NBRC 16594$^{T}$ | 92.7 | 92.77 | 92.77 | 95.72 | 94.78 | 95.18 | 95.86 | 95.85 | 93.58 | 97.03 | 100 | 96.93 | 96.72 |
| *Tanticharoenia sakaeratensis* NBRC 103193$^{T}$ | 92.58 | 92.64 | 92.64 | 95.86 | 93.99 | 95.34 | 95.86 | 96.12 | 93.24 | 98.02 | 96.93 | 100 | 95.86 |
| *Acetobacter aceti* DSM 3508$^{T}$ | 92.57 | 92.63 | 92.63 | 95.18 | 95.11 | 94.61 | 95.92 | 95.85 | 93.97 | 96.24 | 96.72 | 95.86 | 100 |

**Supplementary data Table S6**. Genome annotation and RAST subsystems classification of functionally important genes

| Category | Subsystem | Role | Ha5 | Ta1 | Jb2 |
|---|---|---|---|---|---|
| Cofactors, Vitamins, Prosthetic Groups, Pigments | Biotin biosynthesis | Dethiobiotin synthetase (EC 6.3.3.3) | E3E11_RS04390 | E3203_08300 | E3202_05740 |
| | | Biotin synthase (EC 2.8.1.6) | E3E11_RS00155 | E3203_04135 | E3202_01565 |
| | | Adenosylmethionine-8-amino-7-oxononanoate aminotransferase (EC 2.6.1.62) | E3E11_RS04410 | E3203_08320 | E3202_05760 |
| | | 8-amino-7-oxononanoate synthase (EC 2.3.1.47) | E3E11_RS04405 | E3203_08315 | E3202_05755 |
| | Thiamin biosynthesis | Thiamine-monophosphate kinase (EC 2.7.4.16) | E3E11_RS08340 | E3203_03910 | E3202_01340 |
| | | 1-deoxy-D-xylulose 5-phosphate synthase (EC 2.2.1.7) | E3E11_RS07135 | E3203_02710 | E3202_00110 |
| | Riboflavin, FMN and FAD metabolism | FMN adenylyltransferase (EC 2.7.7.2) | E3E11_RS04780 | E3203_00365 | E3202_06135 |
| | | Riboflavin synthase eubacterial/eukaryotic (EC 2.5.1.9) | E3E11_RS03675 | E3203_07565 | E3202_05050 |
| | | Diaminohydroxyphosphoribosylaminopyrimidine deaminase (EC 3.5.4.26) | E3E11_RS03680 | E3203_07570 | E3202_05055 |
| | | 6,7-dimethyl-8-ribityllumazine synthase (EC 2.5.1.78) | E3E11_RS03665 | E3203_07555 | E3202_05040 |
| | | 5-amino-6-(5-phosphoribosylamino)uracil reductase (EC 1.1.1.193) | E3E11_RS03680 | E3203_07570 | E3202_05055 |
| | | Riboflavin kinase (EC 2.7.1.26) | E3E11_RS04780 | E3203_00365 | E3202_06135 |
| | | GTP cyclohydrolase II (EC 3.5.4.25) | E3E11_RS03670 | E3203_07560 | E3202_05045 |
| | | 3,4-dihydroxy-2-butanone 4-phosphate synthase (EC 4.1.99.12) | E3E11_RS03670 | E3203_07560 | E3202_05045 |
| | Pyridoxin (Vitamin B6) Biosynthesis | D-3-phosphoglycerate dehydrogenase (EC 1.1.1.95) | E3E11_RS07705 | E3203_03275 | E3202_00700 |
| | | 4-hydroxythreonine-4-phosphate dehydrogenase (EC 1.1.1.262) | E3E11_RS08200 | E3203_03770 | E3202_01200 |
| | | Pyridoxamine 5'-phosphate oxidase (EC 1.4.3.5) | E3E11_RS00320 | E3203_04305 | E3202_01735 |
| | | Pyridoxine 5'-phosphate synthase (EC 2.6.99.2) | E3E11_RS08205 | E3203_03775 | E3202_01205 |
| | | Phosphoserine aminotransferase (EC 2.6.1.52) | E3E11_RS05065 | E3203_00650 | E3202_06420 |
| | | NAD-dependent glyceraldehyde-3-phosphate dehydrogenase (EC 1.2.1.12) | E3E11_RS05710 | E3203_01295 | E3202_07065 |
| | | 1-deoxy-D-xylulose 5-phosphate synthase (EC 2.2.1.7) | E3E11_RS07135 | E3203_02710 | E3202_00110 |
| | Folate biosynthesis cluster | GTP cyclohydrolase I (EC 3.5.4.16) type 1 | E3E11_RS03035 | E3203_06950 | E3202_04435 |
| | | Pantoate--beta-alanine ligase (EC 6.3.2.1) | E3E11_RS06465 | E3203_02050 | E3202_07820 |
| | | Dihydropteroate synthase (EC 2.5.1.15) | E3E11_RS03045 | E3203_06960 | E3202_04445 |
| | | Dihydroneopterin aldolase (EC 4.1.2.25) | E3E11_RS03040 | E3203_06955 | E3202_04440 |
| | | 2-amino-4-hydroxy-6-hydroxymethyldihydropteridine pyrophosphokinase (EC 2.7.6.3) | E3E11_RS03040 | E3203_06955 | E3202_04440 |
| | | tRNA(Ile)-lysidine synthetase (EC 6.3.4.19) | E3E11_RS06365 | E3203_01950 | E3202_07720 |
| Stress Response | Oxidative stress | Catalase (EC 1.11.1.6) | E3E11_RS04075 | E3203_07990 | E3202_05440 |
| Amino Acids and Derivatives | Glutamine, Glutamate, Aspartate and Asparagine Biosynthesis | Glutamine synthetase type I (EC 6.3.1.2) | E3E11_RS04145 | E3203_08060 | E3202_05510 |
| | | Aspartate aminotransferase (EC 2.6.1.1) | E3E11_RS04750 | E3203_00335 | E3202_06105 |
| | | Glutamate racemase (EC 5.1.1.3) | E3E11_RS00290 | E3203_04270 | E3202_01700 |
| | Glutamine synthetases | Glutamine synthetase type I (EC 6.3.1.2) | E3E11_RS04145 | E3203_08060 | E3202_05510 |
| | Histidine Biosynthesis | Phosphoribosylformimino-5-aminoimidazole carboxamide ribotide isomerase (EC 5.3.1.16) | E3E11_RS02345 | E3203_06290 | E3202_03765 |
| | | Phosphoribosyl-ATP pyrophosphatase (EC 3.6.1.31) | E3E11_RS02355 | E3203_06300 | E3202_03775 |

| | | | | |
|---|---|---|---|---|
| | Histidinol-phosphatase [alternative form] (EC 3.1.3.15) | E3E11_RS07710 | E3203_03280 | E3202_00705 |
| | Histidinol-phosphate aminotransferase (EC 2.6.1.9) | E3E11_RS06400 | E3203_01985 | E3202_07755 |
| | Imidazoleglycerol-phosphate dehydratase (EC 4.2.1.19) | E3E11_RS02335 | E3203_06280 | E3202_03755 |
| | ATP phosphoribosyltransferase regulatory subunit (EC 2.4.2.17) | E3E11_RS05070 | E3203_00655 | E3202_06425 |
| | Imidazole glycerol phosphate synthase amidotransferase subunit (EC 2.4.2.-) | E3E11_RS02350 | E3203_06295 | E3202_03770 |
| | Phosphoribosyl-AMP cyclohydrolase (EC 3.5.4.19) | E3E11_RS00190 | E3203_04170 | E3202_01600 |
| | Histidinol dehydrogenase (EC 1.1.1.23) | E3E11_RS03615 | E3203_07505 | E3202_04990 |
| Arginine Biosynthesis | N-succinyl-L,L-diaminopimelate desuccinylase (EC 3.5.1.18) | E3E11_RS07225 | E3203_02800 | E3202_00200 |
| | Glutamate N-acetyltransferase (EC 2.3.1.35) | E3E11_RS06295 | E3203_01880 | E3202_07650 |
| | Acetylglutamate kinase (EC 2.7.2.8) | E3E11_RS07210 | E3203_02785 | E3202_00185 |
| | Argininosuccinate lyase (EC 4.3.2.1) | E3E11_RS00435 | E3203_04420 | E3202_01850 |
| | N-acetylglutamate synthase (EC 2.3.1.1) | E3E11_RS06295 | E3203_01880 | E3202_07650 |
| | Argininosuccinate synthase (EC 6.3.4.5) | E3E11_RS01350 | E3203_05330 | E3202_02770 |
| | N-acetyl-gamma-glutamyl-phosphate reductase (EC 1.2.1.38) | E3E11_RS05045 | E3203_00630 | E3202_06400 |
| | Ornithine carbamoyltransferase (EC 2.1.3.3) | E3E11_RS04745 | E3203_00330 | E3202_06100 |
| | Acetylornithine aminotransferase (EC 2.6.1.11) | E3E11_RS04750 | E3203_00335 | E3202_06105 |
| Methionine Biosynthesis | Homoserine dehydrogenase (EC 1.1.1.3) | E3E11_RS03125 | E3203_07040 | E3202_04530 |
| | Methionine ABC transporter substrate-binding protein | E3E11_RS02830 | E3203_06760 | E3202_04240 |
| | Methionine ABC transporter ATP-binding protein | E3E11_RS02835 | E3203_06765 | E3202_04245 |
| | S-adenosylmethionine synthetase (EC 2.5.1.6) | E3E11_RS02600 | E3203_06540 | E3202_04010 |
| | Serine acetyltransferase (EC 2.3.1.30) | E3E11_RS02055 | E3203_06040 | E3202_03480 |
| | Homoserine kinase (EC 2.7.1.39) | E3E11_RS01115 | E3203_05105 | E3202_02535 |
| | Cysteine synthase (EC 2.5.1.47) | E3E11_RS04690 | E3203_00275 | E3202_06045 |
| | Methionine ABC transporter permease protein | E3E11_RS02840 | E3203_06770 | E3202_04250 |
| | Adenosylhomocysteinase (EC 3.3.1.1) | E3E11_RS04065 | E3203_07980 | E3202_05430 |
| Threonine and Homoserine Biosynthesis | Homoserine dehydrogenase (EC 1.1.1.3) | E3E11_RS03125 | E3203_07040 | E3202_04530 |
| | Aspartate-semialdehyde dehydrogenase (EC 1.2.1.11) | E3E11_RS02645 | E3203_06580 | E3202_04050 |
| | Aspartate aminotransferase (EC 2.6.1.1) | E3E11_RS04750 | E3203_00335 | E3202_06105 |
| | Threonine synthase (EC 4.2.3.1) | E3E11_RS07340 | E3203_02915 | E3202_00315 |
| | Homoserine kinase (EC 2.7.1.39) | E3E11_RS01115 | E3203_05105 | E3202_02535 |
| | Aspartokinase (EC 2.7.2.4) | E3E11_RS00555 | E3203_04540 | E3202_01970 |
| Cysteine Biosynthesis | Cysteine synthase (EC 2.5.1.47) | E3E11_RS04690 | E3203_00275 | E3202_06045 |
| | Sulfate permease | E3E11_RS05380 | E3203_00965 | E3202_06735 |
| | Serine acetyltransferase (EC 2.3.1.30) | E3E11_RS02055 | E3203_06040 | E3202_03480 |
| Lysine Biosynthesis DAP Pathway | N-succinyl-L,L-diaminopimelate desuccinylase (EC 3.5.1.18) | E3E11_RS07225 | E3203_02800 | E3202_00200 |
| | 2,3,4,5-tetrahydropyridine-2,6-dicarboxylate N-succinyltransferase (EC 2.3.1.117) | E3E11_RS07220 | E3203_02795 | E3202_00195 |
| | Diaminopimelate epimerase (EC 5.1.1.7) | E3E11_RS05090 | E3203_00675 | E3202_06445 |
| | Aspartate-semialdehyde dehydrogenase (EC 1.2.1.11) | E3E11_RS02645 | E3203_06580 | E3202_04050 |
| | 4-hydroxy-tetrahydrodipicolinate synthase (EC 4.3.3.7) | E3E11_RS07725 | E3203_03295 | E3202_00720 |
| | Diaminopimelate decarboxylase (EC 4.1.1.20) | E3E11_RS07580 | E3203_03150 | E3202_00555 |

| | | | | | |
|---|---|---|---|---|---|
| | | 4-hydroxy-tetrahydrodipicolinate reductase (EC 1.17.1.8) | E3E11_RS02595 | E3203_06535 | E3202_04005 |
| | | Aspartokinase (EC 2.7.2.4) | E3E11_RS00555 | E3203_04540 | E3202_01970 |
| | Chorismate: Intermediate for synthesis of Tryptophan, PAPA antibiotics, PABA, 3-hydroxyanthranilate and more. | Phosphoribosylformimino-5-aminoimidazole carboxamide ribotide isomerase (EC 5.3.1.16) | E3E11_RS02345 | E3203_06290 | E3202_03765 |
| | | Anthranilate synthase, amidotransferase component (EC 4.1.3.27) | E3E11_RS07875 | E3203_03445 | E3202_00875 |
| | | Aminodeoxychorismate lyase (EC 4.1.3.38) | E3E11_RS02905 | E3203_06830 | E3202_04315 |
| | | Tryptophan synthase alpha chain (EC 4.2.1.20) | E3E11_RS02670 | E3203_06605 | E3202_04075 |
| | | Anthranilate phosphoribosyltransferase (EC 2.4.2.18) | E3E11_RS07880 | E3203_03450 | E3202_00880 |
| | | Tryptophan synthase beta chain (EC 4.2.1.20) | E3E11_RS02665 | E3203_06600 | E3202_04070 |
| | | Indole-3-glycerol phosphate synthase (EC 4.1.1.48) | E3E11_RS07885 | E3203_03455 | E3202_00885 |
| | | Anthranilate synthase, aminase component (EC 4.1.3.27) | E3E11_RS07870 | E3203_03440 | E3202_00870 |
| | | Phosphoribosylanthranilate isomerase (EC 5.3.1.24) | E3E11_RS08170 | E3203_03740 | E3202_01170 |
| | | Para-aminobenzoate synthase, aminase component (EC 2.6.1.85) | E3E11_RS02900 | E3203_06825 | E3202_04310 |
| | | Para-aminobenzoate synthase, amidotransferase component (EC 2.6.1.85) | E3E11_RS07875 | E3203_03445 | E3202_00875 |
| | Phenylalanine and Tyrosine Branches from Chorismate | Chorismate mutase I (EC 5.4.99.5) | E3E11_RS00275 | E3203_04255 | E3202_01685 |
| | | Prephenate dehydratase (EC 4.2.1.51) | E3E11_RS07720 | E3203_03290 | E3202_00715 |
| | | Cyclohexadienyl dehydrogenase (EC 1.3.1.12)(EC 1.3.1.43) | E3E11_RS07625 | E3203_03195 | E3202_00620 |
| | Tryptophan synthesis | Anthranilate synthase, amidotransferase component (EC 4.1.3.27) | E3E11_RS07875 | E3203_03445 | E3202_00875 |
| | | Aminodeoxychorismate lyase (EC 4.1.3.38) | E3E11_RS02905 | E3203_06830 | E3202_04315 |
| | | Tryptophan synthase alpha chain (EC 4.2.1.20) | E3E11_RS02670 | E3203_06605 | E3202_04075 |
| | | Para-aminobenzoate synthase, aminase component (EC 2.6.1.85) | E3E11_RS02900 | E3203_06825 | E3202_04310 |
| | | Anthranilate phosphoribosyltransferase (EC 2.4.2.18) | E3E11_RS07880 | E3203_03450 | E3202_00880 |
| | | Tryptophan synthase beta chain (EC 4.2.1.20) | E3E11_RS02665 | E3203_06600 | E3202_04070 |
| | | Indole-3-glycerol phosphate synthase (EC 4.1.1.48) | E3E11_RS07885 | E3203_03455 | E3202_00885 |
| | | Para-aminobenzoate synthase, amidotransferase component (EC 2.6.1.85) | E3E11_RS07875 | E3203_03445 | E3202_00875 |
| | | Anthranilate synthase, aminase component (EC 4.1.3.27) | E3E11_RS07870 | E3203_03440 | E3202_00870 |
| | | Phosphoribosylanthranilate isomerase (EC 5.3.1.24) | E3E11_RS08170 | E3203_03740 | E3202_01170 |
| | Proline Synthesis | Pyrroline-5-carboxylate reductase (EC 1.5.1.2) | E3E11_RS04275 | E3203_08190 | E3202_05635 |
| | Glycine Biosynthesis | Serine hydroxymethyltransferase (EC 2.1.2.1) | E3E11_RS08005 | E3203_03575 | E3202_01005 |
| | Alanine biosynthesis | Branched-chain amino acid aminotransferase (EC 2.6.1.42) | E3E11_RS05415 | E3203_01000 | E3202_06770 |
| | | Ferredoxin, 2Fe-2S | E3E11_RS04585 | E3203_00170 | E3202_05940 |
| | | Cysteine desulfurase (EC 2.8.1.7) | E3E11_RS04580 | E3203_04720 | E3202_05935 |
| | | Alanine racemase (EC 5.1.1.1) | E3E11_RS00050 | E3203_04030 | E3202_01460 |
| | | Iron-sulfur cluster regulator IscR | E3E11_RS00755 | E3203_04740 | E3202_02170 |
| | Serine Biosynthesis | D-3-phosphoglycerate dehydrogenase (EC 1.1.1.95) | E3E11_RS07705 | E3203_03275 | E3202_00700 |
| | | Phosphoserine phosphatase (EC 3.1.3.3) | E3E11_RS04095 | E3203_08010 | E3202_05460 |
| | | Serine hydroxymethyltransferase (EC 2.1.2.1) | E3E11_RS08005 | E3203_03575 | E3202_01005 |
| | | Phosphoserine aminotransferase (EC 2.6.1.52) | E3E11_RS05065 | E3203_00650 | E3202_06420 |
| Respiration | Terminal cytochrome oxidases | Cytochrome d ubiquinol oxidase subunit I (EC 1.10.3.-) | E3E11_RS02810 | E3203_06740 | E3202_04220 |
| | | Cytochrome d ubiquinol oxidase subunit II (EC 1.10.3.-) | E3E11_RS02815 | E3203_06745 | E3202_04225 |
